# The effects of population size histories on estimates of selection coefficients from time-series genetic data

**DOI:** 10.1101/048355

**Authors:** Ethan M. Jewett, Matthias Steinrücken, Yun S. Song

**Author notes:** **Corresponding author:** Yun S. Song.

## Abstract

Many approaches have been developed for inferring selection coefficients from time series data while accounting for genetic drift. However, the improvement in inference accuracy that can be attained by modeling drift is unknown. Here, by comparing maximum likelihood estimates of selection coefficients that account for the true population size history with estimates that ignore drift, we address the following questions: how much can modeling the population size history improve estimates of selection coefficients? How much can mis-inferred population sizes hurt inferences of selection coefficients? We conduct our analysis under the discrete Wright-Fisher model by deriving the exact probability of an allele frequency trajectory in a population of time-varying size and we replicate our results under the diffusion model by extending the exact probability of a frequency trajectory derived by Steinrücken *et al*. (2014) to the case of a piecewise constant population. For both the discrete Wright-Fisher and diffusion models, we find that ignoring drift leads to estimates of selection coefficients that are nearly as accurate as estimates that account for the true population history, even when population sizes are small and drift is high. In populations of time-varying size, estimates of selection coefficients that ignore drift are similar in accuracy to estimates that rely on crude, yet reasonable, estimates of the population history. These results are of interest because inference methods that ignore drift are widely used in evolutionary studies and can be many orders of magnitude faster than methods that account for population sizes.

## 1. Introduction

Methods for inferring the selection coefficient at a single genetic locus from time series data have been employed extensively in evolutionary studies of simple traits. Such methods track the frequency of an allele or Mendelian trait over multiple generations and infer the selection coefficient that best explains the observed frequency changes. Studies of selective pressures conducted using time series approaches have provided evidence for strong selective forces in natural populations and have helped to characterize the ways in which environmental factors influence evolution through selection (Clarke and Murray, 1962; Clark, 1979; Wall *et al*., 1980; Lynch, 1987; Stine and Smith, 1990; Goudsmit *et al*., 1996; Cowie and Jones, 1998; Harrigan *et al*., 1998; Cook *et al*., 1999; Haubruge and Arnaud, 2001; Bonhoeffer *et al*., 2002; Reimchen and Nosil, 2002; Cook *et al*., 2005; Labbé *et al*., 2009).

Because random fluctuations in allele frequencies due to genetic drift are often small compared to changes due to selective pressures, it is common practice for studies to assume that allele frequencies change deterministically over time according to well-known deterministic formulas of Fisher (1922, p.424) and Haldane (1927, p.840) or related expressions (Gillespie, 2010; Hartl and Clark, 2007). However, because allele frequency trajectories can be heavily influenced by genetic drift when population sizes or selection coefficients are small, many methods have been developed to account for drift by explicitly modeling finite population sizes when inferring selection coefficients from observed allele frequency trajectories (Manly, 1985; O'Hara, 2005; Bollback *et al*., 2008; Malaspinas *et al*., 2012; Mathieson and McVean, 2013; Lacerda and Seoighe, 2014; Steinrücken *et al*., 2014; Foll *et al*., 2015; Ferrer-Admetlla *et al*., 2015) and when testing hypotheses about selection versus drift (Fisher and Ford, 1947; Schaffer *et al*., 1977; Wilson, 1980; Nishino, 2013; Feder *et al*., 2014; Topa *et al*., 2015).

Although estimates of selection coefficients are likely to be improved by accounting for population size histories, the expected amount of improvement is not well characterized. Even in relatively small populations, allele frequencies and other evolutionary processes behave almost deterministically if the selection coefficient or allele frequency is sufficiently high (Rouzine *et al*., 2001), suggesting that methods that ignore drift might perform well under these conditions. Conversely, if drift is strong allele frequency trajectories can be noisy and the accuracy of methods that ignore drift may be comparable to that of methods that account for population size, as all methods are likely to perform poorly under these conditions (Gallet *et al*., 2012).

If computationally fast methods that ignore drift are accurate, they could dramatically reduce the time required to infer selection coefficients in data sets with many loci. In addition to their computational efficiency, methods that ignore drift do not require estimates of effective population sizes, which can be difficult to obtain accurately. Moreover, ignoring drift can lead to simple formulas and inference procedures under complicated evolutionary scenarios (e.g., Illingworth *et al*., 2012). Therefore, in light of the beneficial properties of methods that ignore drift and assume deterministic allele frequency trajectories, it is of interest to compare their accuracy to that of methods that account for population size histories.

The theoretical accuracy of methods for inferring selection coefficients can be difficult to derive analytically. Thus, to explore differences between methods that ignore or account for drift, one can take the approach of empirically comparing inferences made by the same estimator, either accounting for the true population size history or ignoring the size history by assuming that populations are large and drift is negligible. This is the the approach we take here. For our analyses, we consider maximum likelihood estimators of selection coefficients because they are typically quite accurate and have desirable statistical properties. Moreover, the majority of recently-developed methods for inferring selection coefficients from time series data are maximum likelihood estimators, making them an important category of methods to evaluate.

To draw conclusions about the accuracy of maximum likelihood estimators, it is important to consider estimators based on exact likelihoods rather than approximations, so that differences in estimates can be attributed entirely to whether a method ignores or accounts for drift. Although several approximate approaches have been developed for computing the likelihood of a selection model given time series allele frequency data, only two existing methods compute probabilities that are exact under a widely accepted model. In particular, the methods of Bollback *et al*. (2008) and Steinrücken *et al*. (2014) compute exact probabilities under the diffusion approximation of the Wright-Fisher process. However, no method computes the exact probability of an allele frequency trajectory under the discrete Wright-Fisher model, as the matrix powers required for such a method are considered to be computationally inefficient. Moreover, no existing inference method based on the exact likelihood models time-varying population histories, making it difficult to explore the effects of accounting for demography on inference accuracy.

Here, we derive the exact probability of an allele frequency trajectory in a population of piecewise constant size under two classical models: the discrete Wright-Fisher model and the diffusion approximation of the Wright-Fisher process.

We then use maximum likelihood estimators obtained using these probabilities to explore how ignoring or accounting for the true population history affects estimates of selection coefficients.

## Results

To compare the performance of estimators that ignore or account for drift, we inferred selection coefficients from allele frequency trajectories simulated under a variety of population histories of time-varying size.

### 1.1. The population model.

In all of our analyses, we considered a single biallelic locus with alleles labeled *a* and *A* evolving under selection and recurrent mutation in a panmictic population comprised of *L* different epochs *ℓ* = 1,…*L*, each with constant size *N*_*ℓ*_ diploid individuals (Figure 1). Epoch *ℓ* corresponds to the time interval [*τ*_*ℓ*–1_, *τ*_*ℓ*_], where time is measured continuously in units of generations and we define *τ*_0_ ≡ 0. By varying the population sizes *N*_*ℓ*_ across epochs, it is possible to model a variety of size-change patterns including exponential growth, bottlenecks, and rapidly oscillating population sizes.

**Figure 1.**
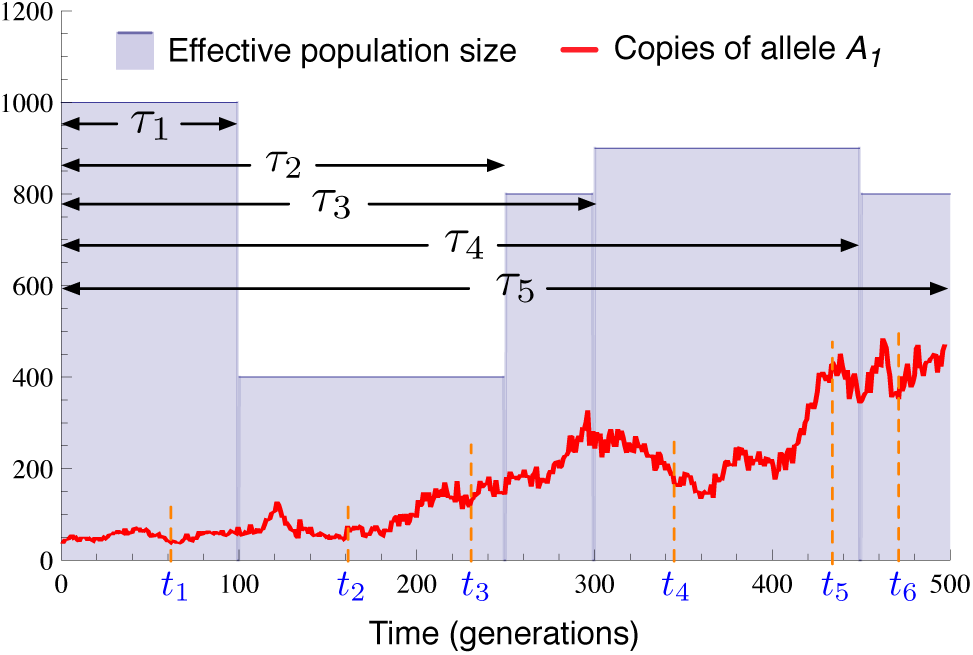
Diagram of the model. An allele at a single locus evolves in a population of piecewise constant size with *L* = 5 epochs spanning the time periods [*τ*_0_,*τ*_1_],…,[*τ*_*L*–1_,*τ*_*L*_], where *τ*_0_ = 0. Samples of sizes *n*_1_,…, *n*_*k*_ haplotypes are taken at times *t*_1_,…,*t*_*K*_.

Within epoch *ℓ*, all mutation and selection parameters are assumed to be constant. In particular, we assume that the per-generation probability that allele *a* mutates to allele *A* is 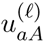 and the per-generation probability that allele *A* to *a* is 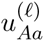 The three possible genotypes, *aa*, *aA*, and *AA*, have relative fitnesses given by 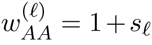, 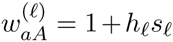, and 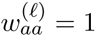 in epoch *ℓ*, where s_o_ is the selection coefficient and ho is the dominance parameter.

We denote the collection of model parameters in epoch *ℓ* by Θ_*ℓ*_ and the set of parameters across all epochs by Θ. It will also be convenient to denote the value of the model parameters at time *t* by 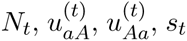, and *h*_t_, where *t* is measured continuously in units of generations. The epoch in which time *t* lies will be denoted by *ℓ*_*t*_ and the epoch in which sampling event *k* lies will be denoted by *ℓ*_*k*_. It will be clear from the context whether the subscript on *ℓ* refers to a time or a sampling event.

We denote the population-wide number of copies of allele *A* in generation *t* by *c*_*t*_ and the population-wide frequency of allele *A* by *yt*. In practice, we do not observe the true population count of allele A· Instead, the data consist of observed counts *o*_1_,…,*o*_*K*_ of the number of times allele *A* is observed in *K* different samples of sizes *n*_1_,…, *n*_*K*_ haplotypes, taken at times *t*_1_ < … < *t*_*K*_. For simplicity, we assume that each sampling time *t*_*k*_ is an integer for *k* = 1,…, *K*. The consecutive observed counts (*o*_*k*_, *o*_*k*+1_,…, *o*_*k′*_) will be denoted by *o*_[*k*:*k′*]_.

In general, we will denote random variables corresponding to observed quantities using capital letters (e.g., *O*_*k*_, *C*_*t*_, and *Y*_*t*_). The goal is to compute the probability ℙ_Θ_{*O*_[1:*K*]_ = *o*_[1:*K*]_} of the observed data, conditional on the model parameters Θ.

### 1.2. Probabilities of frequency trajectories.

Several different evolutionary models can be used to describe stochastic allele frequency changes over time in a population. Discrete changes in allele frequency are often modeled using the Wright-Fisher and Moran processes, whereas continuous changes are often modeled using the diffusion approximation of the Wright-Fisher process (Karlin and Taylor, 1981; Ewens, 2004; Wakeley, 2008) or one of several approximations of the diffusion (e.g. Feder *et al*., 2014; Lacerda and Seoighe, 2014).

Because it is unclear which model provides the most accurate description of biological evolutionary processes, we take the approach in this paper of deriving exact probabilities of allele frequency trajectories under two different evolutionary models: the discrete Wright-Fisher process and the continuous diffusion approximation.

Under the Wright-Fisher model, the probability ℙ_Θ, ***D***_{*O*_[1:*K*]_ = *o*_[1:*K*]_} of the observed allele counts can be obtained using the recursive formula developed in Section 3.1 (Procedure 1). Under the diffusion approximation, the probability ℙ_Θ, ***W***_{*O*_[1:*K*]_ = *o*_[1:*K*]_} can be obtained using the recursive formula developed in Section 3.2 (Procedure 2).

In Sections 3.4.1 and 3.4.2, we show that if drift is ignored and allele frequencies evolve deterministically, then the probabilities ℙ_Θ, ***W***_{*O*_[1:*K*]_ = *o*_[1:*K*]_} and ℙ_Θ, ***D***_{*O*_[1:*K*]_ = *o*_[1:*K*]_} can be reduced to the simpler approximate probabilities 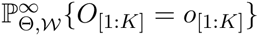 and 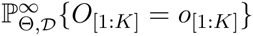 which ignore the population history and which are computed using Procedures 3 and 4, respectively.

Different estimates of the model parameters *Θ* can be obtained using each of the different probabilities ℙ_Θ, ***W***_{*O*_[1:*K*]_ = *o*_[1:*K*]_}, ℙ_Θ, ***D***_{*O*_[1:*K*]_ = *o*_[1:*K*]_}, 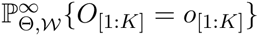 and 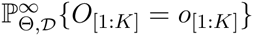 by finding the value of *Θ* that maximizes the given probability of the observed allele counts *o*_[1:*K*]_. In our analyses we estimated the model parameters Θ separately using each of the different probabilities, yielding the estimators *ŝ*_***W***_, *ŝ*_***D***_, 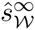, and 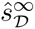. The estimator *ŝ*_***W***_ accounts for drift under the discrete Wright-Fisher model, while drift in this model is ignored by the estimator 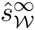. Similarly, the estimator sDaccounts for drift under the diffusion model, while drift in this model is ignored by the estimator 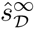.

The degree to which accounting for drift can improve estimates of selection coefficients can be investigated by comparing *ŝ*_***W***_ to 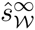 on trajectories simulated under the discrete Wright-Fisher model and by comparing *ŝ*_***D***_ to 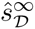 trajectories simulated under the diffusion approximation.

### 1.3. Overview of the experimental design.

We simulated allele frequency trajectories under a variety of selection strengths and piecewise constant population histories reflecting demographic patterns such as exponential growth, bottlenecks, rapid population size oscillations, and constant histories. We then compared the demography-aware estimates *ŝ*_***W***_ and *ŝ*_***D***_ with the estimates 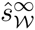 and 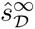 that ignore drift to study the degree to which accounting for population size can improve the accuracy of inferences.

### 1.4. Expected allele frequency trajectories.

Before comparing the accuracy of the different estimators, we first explored the degree to which trajectories that ignore drift differ from trajectories that account for the population size under different evolutionary scenarios. Figure 2 shows the expected frequency of allele A in a discrete Wright-Fisher population of constant size for several different initial allele frequencies, selection coefficients, and effective population sizes. Figure 2 illustrates that, for any starting frequency and selection coefficient, the mean allele frequency trajectory approaches the mean trajectory in a population without drift (e.g., in a population of infinite size), as the true population size increases. Moreover, if the initial frequency is sufficiently high, the expected trajectory is close to its deterministic limit even when the population size is small and drift is high.

**Figure 2.**
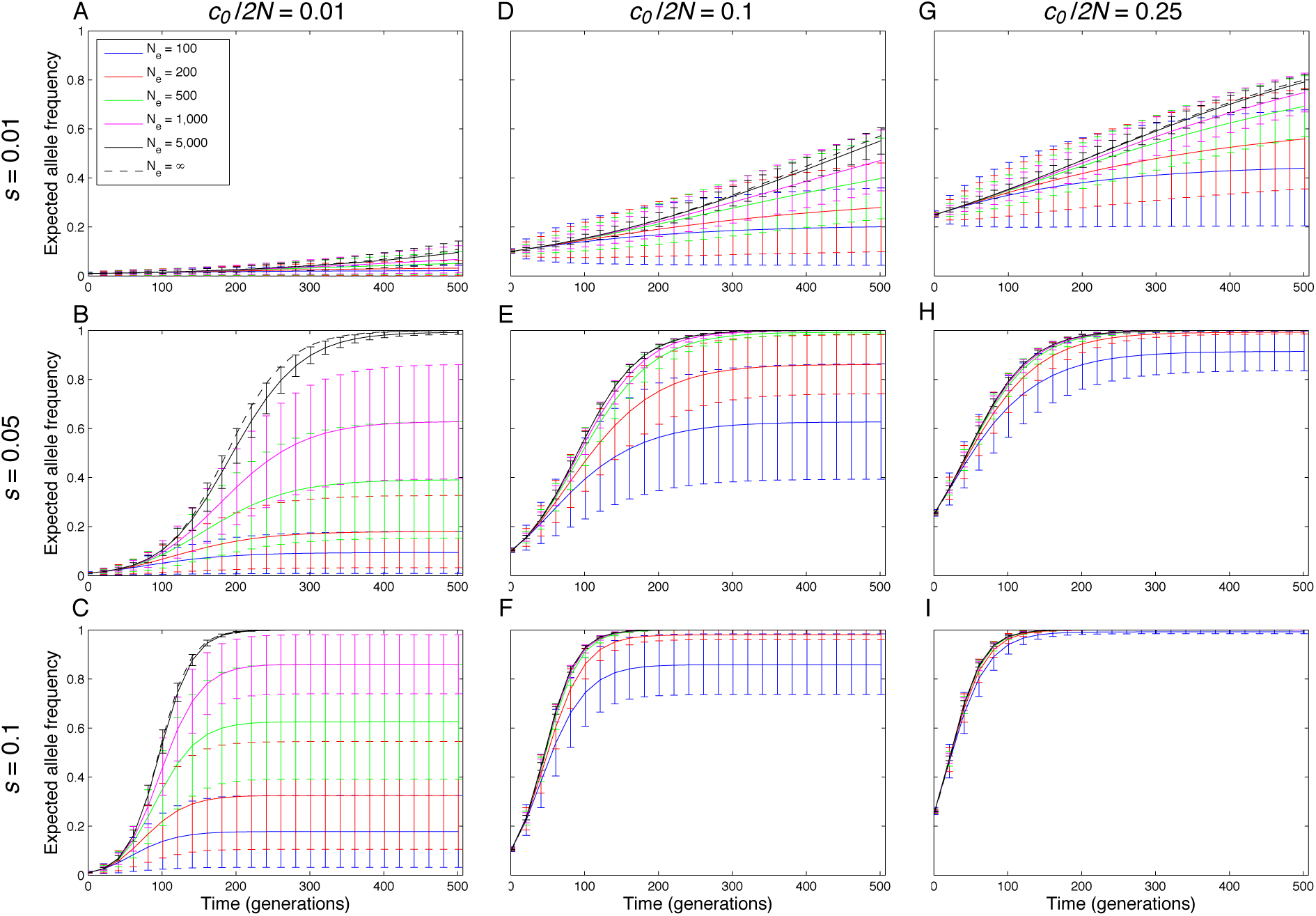
Expected Wright-Fisher trajectories of allele *A* for different initial starting counts *c*_0_, selection coefficients *s*, and effective population sizes *N*. Columns correspond to different initial starting frequencies *c*_0_/2*N* with *c*_0_/2*N* = 0.01, 0.1, and 0.25. In all panels, the error bars show the mean deviation on either side of the expected trajectory (**E**[max{0, *c*_*t*_ ′ **E**[*c*_*t*_]}]/2*N* for the upper bar and **E**[max{0, **E**[*c*_*t*_] – *c*_*t*_}]/2*N* for the lower bar). The dominance parameter is set to *h* = 1/2 in all panels. Because the effects of mutation are negligible during the time periods we consider, we set *u*_*Aa*_ = *u*_*aA*_ = 0.

The results presented in Figure 2 suggest that the allele frequency trajectory will differ substantially from the limiting trajectory without drift only when at least two of the three factors that influence stochasticity in the allele frequency trajectory (effective population size, selection coefficient, and initial allele frequency) are small. Moreover, for biological populations with sufficiently large effective sizes, the allele frequency trajectory is likely to match the deterministic trajectory, regardless of the selection coefficient and initial frequency.

From Figure 2 it can also be seen that an effective population size of several thousand individuals is often sufficiently large to guarantee deterministic behavior, even when the selection coefficient and initial allele frequency are small. Thus, selection coefficient inference methods that ignore drift are likely to be accurate for a broad range of population sizes and selection coefficients. As we will see, methods that ignore drift can be almost as accurate as methods that account for drift, even within the small-parameter-value regime.

### 1.5. Inference accuracy, accounting for constant population size.

To explore how accounting for drift affects inference accuracy, we first considered the accuracy of inferring selection coefficients in a population of constant finite size. Figure 3 shows the maximum likelihood estimate (MLE) of the selection coefficient for three different effective population sizes (*N* = 100, 500, 1000), three selection coefficients (*S* = 0.01, 0.05, 0.1), and two initial allele frequencies (*y*_0_ = 0.01, 0.1) for *h* = 1/2. In each panel, the violin plots summarize the maximum likelihood estimates for 100 different simulation replicates in which an allele frequency trajectory was simulated for 500 generations with samples of size *n* = 50 taken at generations *t* = 50, 100, 150, 200, 250, 300, 350, 400, 450, and 500.

**Figure 3.**
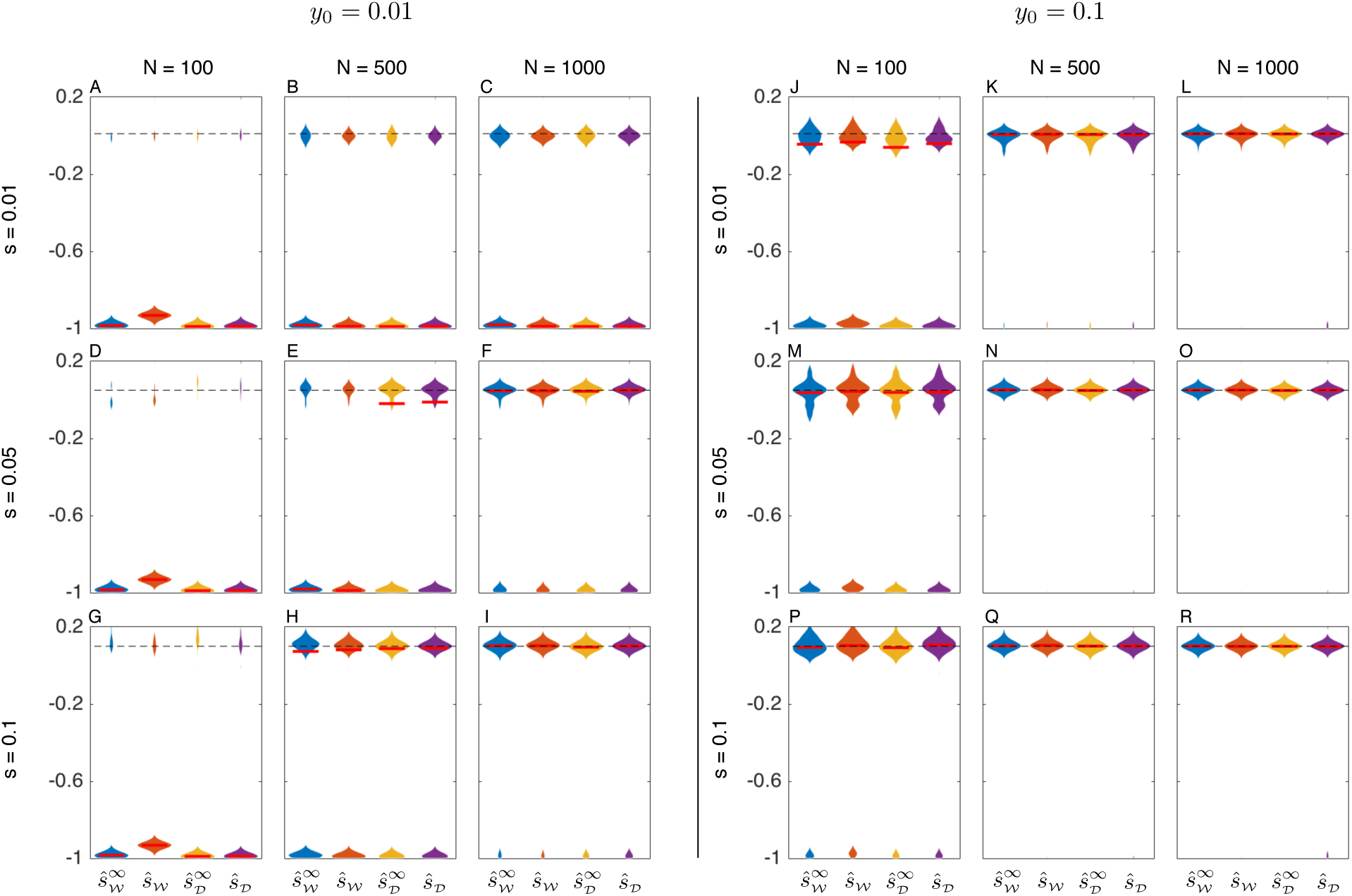
Maximum likelihood estimates of the selection coefficient S in populations of constant size. For each of three different selection coefficients (*s* = 0.01, 0.05, 0.1) and effective population sizes (*N* = 100, 500, 1000), 100 allele frequency trajectories were simulated for 500 generations under the either the Wright-Fisher or diffusion models. Samples of 50 alleles were taken at times 50, 100, 150, 200, 250, 300, 350, 400, 450, and 500 generations. Bimodal violin plots are due to the fact that allele frequency trajectories typically fall into one of two categories: trajectories in which allele *A* is lost quickly, resulting in a strong negative estimate of the selection coefficient, and trajectories in which allele *A* remains segregating long enough to allow a more accurate estimate of the selection coefficient. Red bars indicate medians. The maximum width of each violin plot is scaled to the same value for all estimators.

#### Procedure 1

Computing ℙ_Θ, ***W***_{*O*_[1:*K*]_ = *o*_[1:*K*]_}

1: Define the quantities *d*_0_= (ℙ{*C*_0_ = 0}, ℙ{*C*_0_ = 1},…, ℙ{*C*_0_ = 2N_*t*_0__}) and *γ*(*o*_1_), where *γ*(*o*_*k*_) = (*γ*_0_(*o*_*k*_), *γ*_1_(*o*_*k*_),…, *γ*_2_*N*_*t*_*k*__(*o*_*k*_)) with 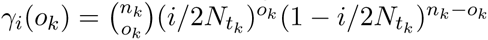.
2: Initialize 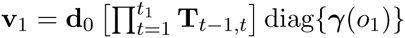.
3: For *k* = 2: *K*, compute

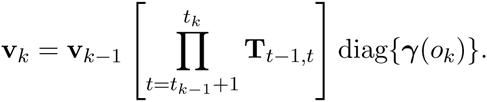
4: Compute 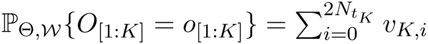

#### Procedure 2

Computing ℙ_Θ, ***D***_{*O*_[1:*K*]_}

1: For an initial starting frequency *x* initialize

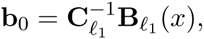

where **B**_*ℓ*_(*x*) is the vector of eigenfunctions of the diffusion operator given in Equation (A.14) and 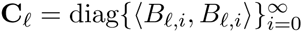 is given in Equation (A.18).
2: For *k* = 1: *K*, compute

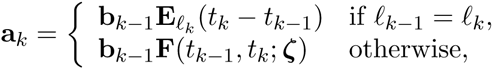

and

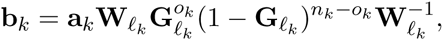

where the matrices **E**_*ℓ*_(*t*), **F**(*t*_*k*–1_, *t*_*k*_; *ζ*), **W**_*ℓ*_, and **G**_*ℓ*_ are given by Equations (A.17), (B.10), (A.15) and (A.11), respectively and *ζ* is the set of Chebyshev nodes in the interval [0,1]. The matrix inverse 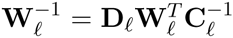 is computed easily using the diagonal matrices **C**_*ℓ*_ and **D**_*ℓ*_ in Equations (A.18) and (A.19).
3: Compute

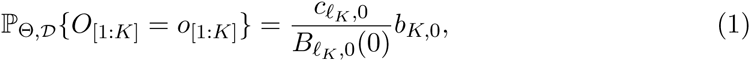

where *c*_*ℓ*_*K*_, _0__ = 〈*B*_*ℓ*, 0_, *B*_*ℓ*, 0_〉 = [*C*_*ℓ*_*K*__]_0, 0_ is the (0,0) element of matrix **C**_*ℓ*_ in Equation (A.18) and *B*_*ℓ*_*K*_, 0_(0) is the 0th element of the vector *B*_*ℓ*_*K*__(0) in Equation (A.14).

For the discrete Wright-Fisher model, allele frequency trajectories were simulated by sampling the allele frequency in each generation from the vector of transition probabilities, conditional on the frequency in the previous generation. Under the diffusion model, trajectories were sampled using the approach of Jenkins and Spanò (2015, personal communication). Maximum likelihood estimates were obtained for the Wright-Fisher trajectories using a grid search over the likelihoods computed using Procedures 1 and 3, and maximum likelihood estimates for the diffusion trajectories were obtained using the same grid search approach over the likelihoods computed using Procedures 2 and 4.

#### Procedure 2

Computing 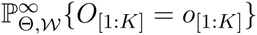

1: Starting with 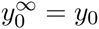, for *t* = 0,…, *t*_*K*_ – 1,

(1) Compute 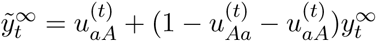
(2) Compute

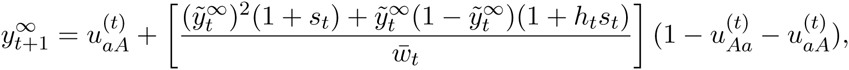

where 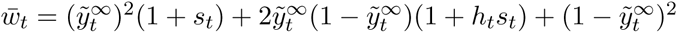.
2: Compute

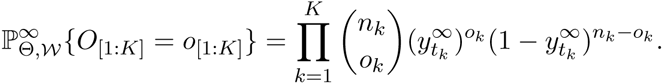

#### Procedure 4

Computing 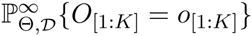

1: Fix a large integer *n* and set Δ*t* = 1/*n*.
2: Starting with 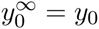, for *j* = 0,…, *nt*_*K*_ – 1, compute

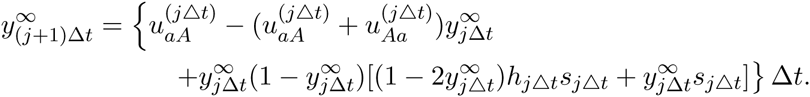
3: Compute

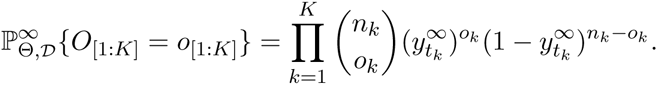

By comparing the estimates computed accounting for drift with the estimates obtained ignoring drift, it can be seen that all methods have similar accuracies. All methods perform well when the population size, selection coefficient, and initial frequency are sufficiently large (e.g., Figure 3I for the case *y*_0_ = 0.01 and Panels 3K through 3R for the case *y*_0_ = 0.1), and all methods perform poorly, otherwise. Figure 3 suggests that the parameter range in which selection coefficients can be inferred accurately by maximum likelihood corresponds with the range in which the assumption *N* ≈ ∞ yields accurate inferences. Put another way: the regime in which selective pressures are strong enough to measure accurately corresponds to the regime in which allele frequency change is quasi-deterministic. Thus, methods that ignore or account for drift are likely to produce estimates of similar accuracy.

### 1.6. Inference accuracy in populations of piecewise constant size.

We next explored the degree to which accounting for more complicated population histories can improve maximum likelihood estimates, focusing on three scenarios, a population with a bottleneck, a population undergoing exponential growth, and a population undergoing rapid oscillations in size. Under each scenario, we simulated 100 allele frequency trajectories for an allele with selection coefficient *s* = 0.05, dominance parameter *h* = 1/2, and initial frequency *y*_0_ = 0. 1 under either the Wright-Fisher or diffusion models. The parameter values in these simulations were chosen so that drift would be strong enough to affect allele frequency trajectories, but not strong enough to prevent accurate inferences of selection coefficients.

To investigate the effect on accuracy of using crude, yet reasonable estimates of the population history, we also inferred selection coefficients using likelihoods computed using variants of Procedures 1 and 2 in which the population was assumed to consist of a single epoch of constant size equal to the Watterson estimate (Hein *et al.*, 2005, p.62) of the effective population size. The Watterson estimate was obtained by computing the expected site frequency spectrum (SFS) for the multi-epoch model for a sample size of 20 alleles, and then inferring the effective size of a single epoch using Watterson’s estimate. The discrete Wright-Fisher and diffusion estimators based on the Watterson estimate of effective size are denoted by 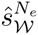 and 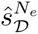, respectively.

#### 1.6.1. The case of a bottleneck.

To model populations with bottlenecks, we considered populations composed of three epochs, each of length 100 generations, with sizes *N*_1_, *N*_2_, and *N*_3_ satisfying *N*_1_ = *N*_3_ = 5*N*_2_. Samples of size 50 were taken at times 50, 100, 150, 200, 250, and 300. Figures 4A and 4B show results for two different populations; in the population in Figure 4A, we set *N*_1_ = 500 and in the population in Figure 4B we set *N*_1_ = 2500.

**Figure 4.**
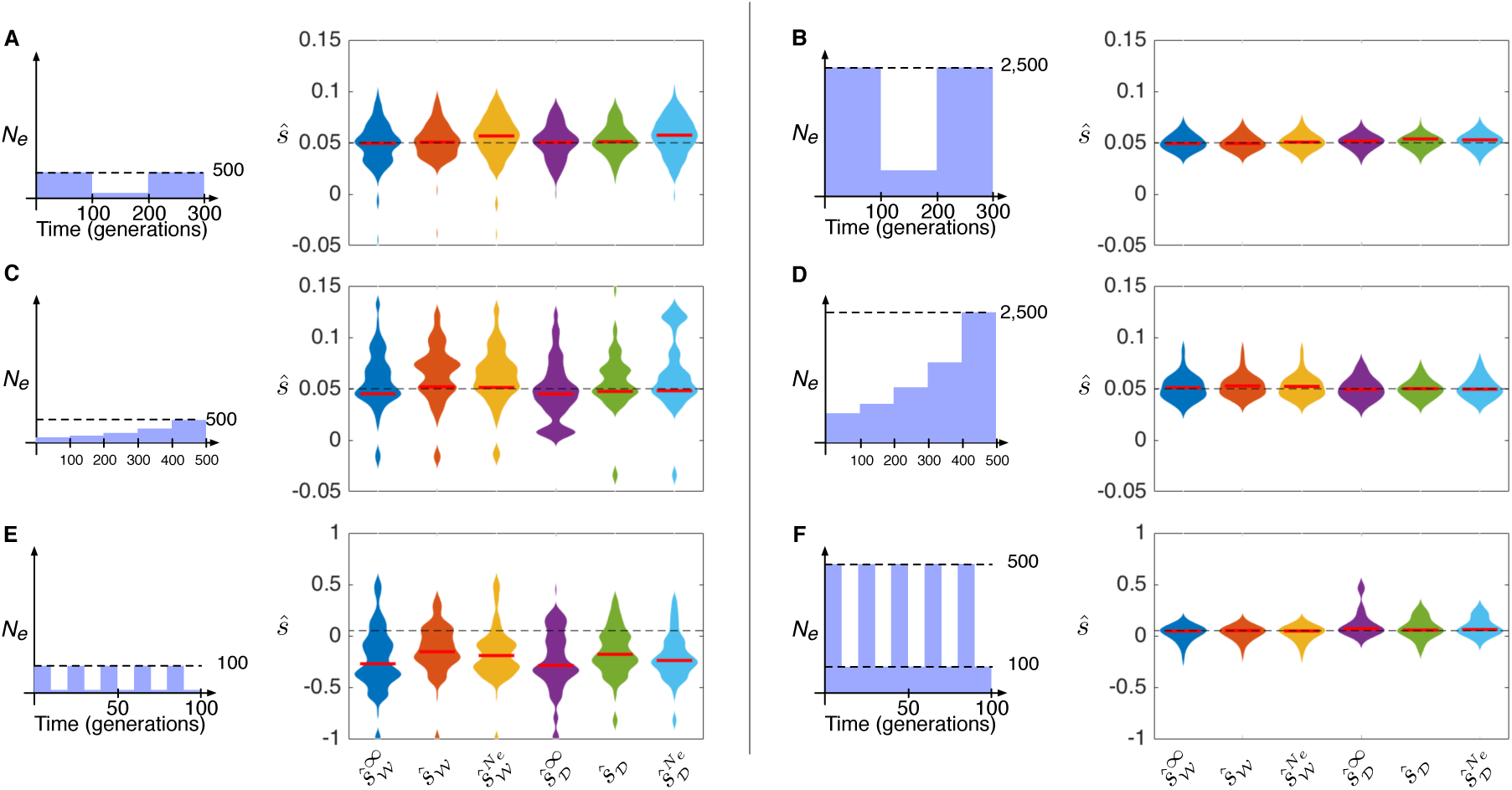
Maximum likelihood estimates of the selection coefficient s in populations with a bottleneck, exponential growth, or rapidly oscillating population size. In each panel, the trajectory of an allele with selection coefficient *s* = 0.05, dominance parameter *h* = 1/2, and starting frequency *y*_0_ = 0.1 was simulated 100 times under the Wright-Fisher and diffusion models. Red bars indicate medians. The maximum width of each violin plot is scaled to the same value for all estimators.

From Figures 4A and 4B, it can be seen that all methods performed similarly. However, the methods that relied on the Watterson estimate of the effective population size were more biased than the other two methods when the effective population size was small, suggesting that methods that ignore drift entirely can produce more accurate estimates than methods that rely on rough estimates of the population history for the bottleneck model. Note that, despite the tight bottleneck in Figure 4A, inferences were still relatively accurate due to the larger sizes of epochs 1 and 3.

#### 1.6.2. The case of exponential growth.

To model exponential growth, we considered populations composed of five epochs, each of length 100 generations, with effective population sizes chosen to represent five-fold exponential growth across all five epochs. Specifically, the size in epoch *ℓ* was set to *N*_*ℓ*_ = *N*_1_*e*^‒*ητ*_*ℓ*–1_^, where the growth constant *η* was chosen such that *e*^‒*ητ*_5_^ = 1/5.

Samples of size 50 were taken in generations 100, 200, 300, 400, and 500. From the results in Figures 4C and 4D, it can be seen that all methods performed with similar accuracy in the growth scenario.

#### 1.6.3. The case of rapidly oscillating population size.

Figures 4E and 4F show inferences of the selection coefficient in a population with rapidly oscillating size. Such demographic histories, which are often seen in insect populations like *Drosophila*, have moderate arithmetic mean sizes, but small harmonic mean sizes and experience episodes of extreme drift.

In the simulations shown in Figure 4E, the population size oscillates rapidly between 10 and 100 diploids every five generations. In the simulations shown in Figure 4F, the population size oscillates between 100 and 500 diploids every five generations. From Figure 4 it can be seen that the methods that ignore drift have similar accuracy to the methods that account for drift, although the methods that account for drift are slightly less biased when the population size oscillates between very small values (Figure 4E).

### 1.7. Conditioning on segregation in the final sample.

It is sometimes of interest to infer the selection coefficient of an allele, conditional on the event that the allele is segregating in the most recent sample. Such conditional inferences are useful if alleles are ascertained in present-day samples and their historical trajectories are subsequently investigated.

Conditioning on segregation in the final sample is also useful for estimating weak positive selection coefficients when initial allele frequencies are low. This is because a large fraction of weakly selected alleles with low initial frequencies will drift out of the population quickly resulting in large negative estimates of their selection coefficients. However, more accurate estimates can be obtained for the subset of alleles that are not lost quickly, which can be seen in Figures 3B, 3C, and 3E, in which the part of the density corresponding to alleles that are not lost quickly from the population is localized around the true selection coefficient.

Considering only alleles that are segregating in the final sample can lead to biased estimates of selection coefficients if likelihood methods do not properly condition on segregation. For example, weakly selected alleles typically drift out of small populations quickly. Thus, weakly selected alleles that escape loss and ultimately fix generally exhibit faster-than-expected increases in frequency that are similar to the unconditional trajectories of alleles under stronger selection. Thus, if a likelihood method does not properly account for conditioning, weakly selected alleles that are segregating in the final sample will have inflated inferred selection coefficients.

Estimators that ignore drift cannot be modified to condition on the event of segregation in the final sample because they implicitly assume that alleles follow fixed trajectories whose long-term behavior in the absence of mutation is entirely determined by the selection coefficient: fixation for positively selected alleles and loss for negatively selected alleles. Thus, estimators that ignore drift are expected to produce biased estimates of selection coefficients when applied to conditioned trajectories.

In contrast, the allele frequency trajectories in likelihood methods that account for the population size are modeled stochastically, allowing likelihoods to be modified to condition on segregation in the final sample. It is expected that methods that account for the true population size can be modified to produce accurate estimates of selection coefficients, whereas methods that ignore drift will necessarily produce biased estimates.

#### 1.7.1. Simulations conditioning on segregation.

To investigate the degree to which accounting for drift can improve estimates of selection coefficients when allele frequency trajectories are conditioned on segregation in the final sample, we modified the discrete Wright-Fisher probability in Section 1 to compute the likelihood conditional on segregation in the final sample using results derived in Section 3.3. Under a grid search, this modified likelihood yields the conditional maximum likelihood estimator *ŝ*_***W***_|_*S*_*K*__. We compared the estimates computed using the exact conditional estimator *ŝ*_***W***_|_*S*_*K*__ with estimates computed using the approximate estimator 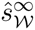 that ignores drift and cannot be modified to account for conditional allele frequency trajectories.

The effect of failing to account for conditioning is evident in the blue violin plots in Figure 5A-I, which correspond to the unconditional approximate maximum likelihood estimates 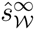. As expected, when the true selection coefficient is small (*s* ≤ 0.01), the estimates 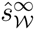 are biased upward. Conversely, when the selection coefficient is larger (*s* ≥ 0.05), the approximate estimator 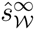 produces negatively biased estimates because alleles under strong positive selection that remain segregating in the final sample show slower-than-expected increases in frequency. In contrast to the estimator 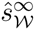, the bias is negligible in the estimator *ŝ*_***W***_|_*S*_*K*__, which accounts for drift and properly conditions on segregation in the final sample (orange violin plots).

**Figure 5.**
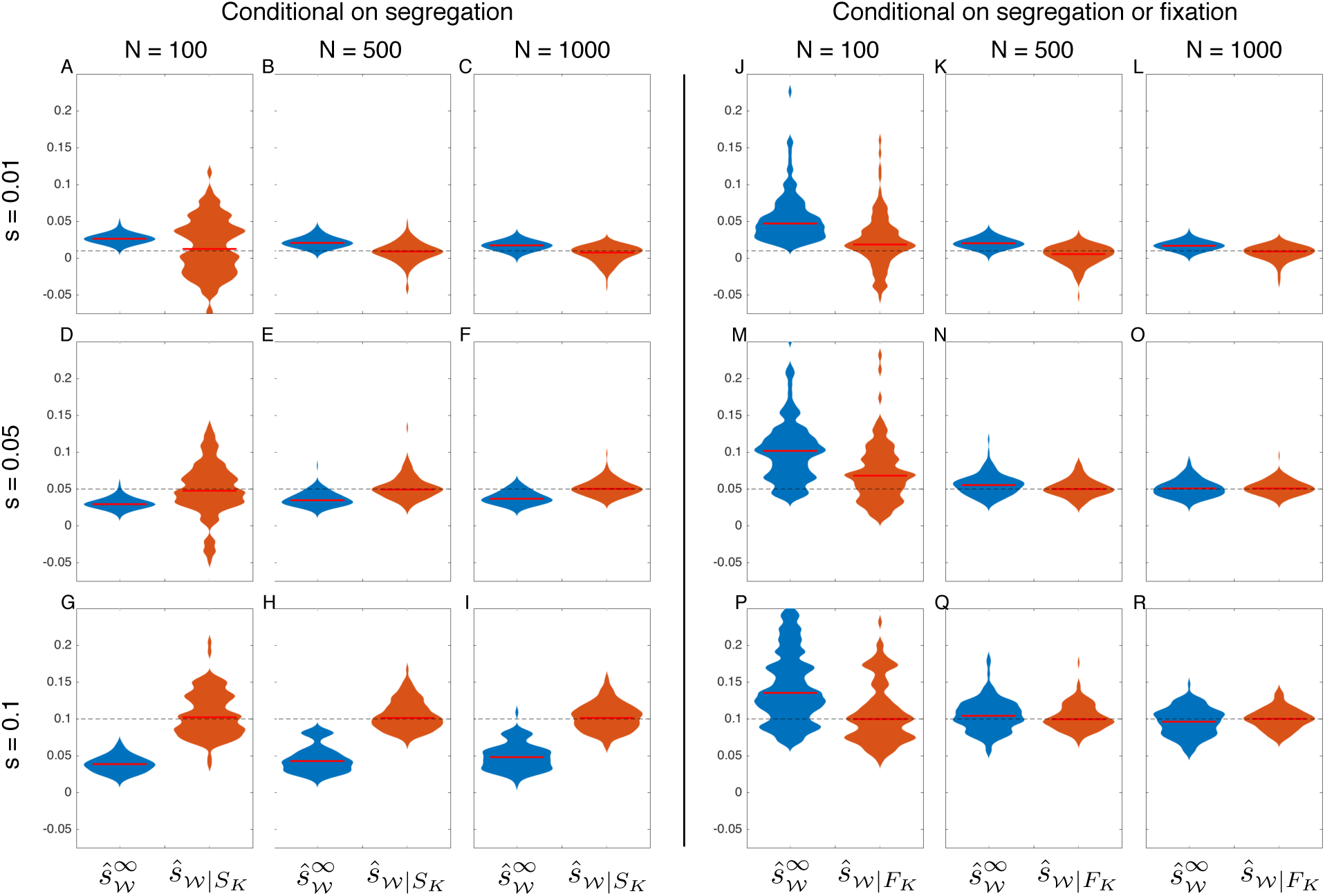
Estimates of selection coefficients, conditional on segregation. Each violin plot was computed using 100 frequency trajectories sampled over 500 generations for an allele with selection coefficient *s* = 0.01 and initial frequency *y*_0_ = 0.01. As in Figure 3, samples of size *n* = 50 were taken in generations 50, 100, 150, 200, 250, 300, 350, 400, 450, and 500. In Panels A-I, trajectories were sampled conditional on the event that the selected allele was segregating in the final sample. In Panels J-R, trajectories were sampled conditional on the event that the selected allele was either segregating or fixed in the final sample. Red bars indicate medians. The maximum width of each violin plot is scaled to the same value for both estimators.

The results shown in Figure 5A-I suggest that methods that account for drift are capable of significantly improving the accuracy of estimates of selection coefficients when allele frequency trajectories are conditioned on segregation. The differences in accuracy between methods that ignore or account for drift are visible for a range of selection coefficients and population sizes. However, the differences in accuracy between the methods diminish as the selection coefficient becomes weaker or the population size becomes larger.

#### 1.7.2 Simulations conditioning on segregation or fixation.

The magnitude of the bias in the estimates 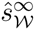 is due in part to the event on which trajectories are conditioned. In cases involving positive selection in populations of moderate or large size, most alleles will be fixed in the final sample (e.g., > 80% fixation within 10 generations when *s* = 0.1, *h* = 1/2, and *y*_0_ = 0. 01, and *N* = 1000). Thus, it may sometimes be more natural to condition on the event *F*_*K*_ that a selected allele is found (segregating or fixed) in the final sample. Under this conditioning scheme, the approximate estimator 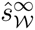 will not generally produce negatively biased estimates of selection coefficients because allele frequency trajectories will not be constrained to those which exhibit slower-than-expected increases in allele frequency.

In light of these considerations, we repeated the analysis shown in Figure 5A-I, simulating allele frequency trajectories conditional on the event that the allele was segregating or fixed in the final sample. To compare the estimates 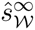 with maximum likelihood estimates that fully account for drift and the proper conditioning, we also modified the probability in ℙ_Θ, ***W***_{*O*_[1:*K*]_ = *o*_[1:*K*]_} computed in Procedure 1 to condition on the event *F*_*K*_ of segregation or fixation in the final sample, yielding the conditional probability ℙ_Θ, ***D***_{*O*_[1:*K*]_ = *o*_[1:*K*]_|*F*_*K*_} (Equation 19) with the associated estimator *ŝ*_***W***_|_*F*_*K*__.

By comparing Figure 5J-R with Figure 5A-I, it can be seen that the estimator 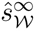 has considerably less bias when conditioning on the event *F*_*K*_ than when conditioning on *S*_*K*_. Although the bias is still high when the population size is small (*N* ≈ 100), it decreases quickly as the population size increases and becomes comparable to the bias in the properly conditioned, demography-aware estimator *ŝ*_***W***_|_*F*_*K*__ when the population size is greater than approximately *N* = 500 diploids. In contrast to Figure 5E-I, the bias in 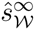 observed in Figure 5M-R is positive because the trajectories on which these estimates are based exclude those in which the allele is lost; thus, they exhibit faster-than-expected growth on average.

The results in Figure 5J-R suggest that under certain conditioning schemes, methods that ignore drift can produce similar estimates to methods that account for drift.

### 1.8. The effect of sample size on accuracy.

When the sample size is small, the variance in estimates arising from sampling noise will tend to obscure small differences between estimators that ignore or account for population size. Thus, when comparing methods, it is important to sample a sufficiently large number of alleles to ensure that the differences between the methods due to ignoring or accounting for drift are visible.

To evaluate the effects of sample size on inference accuracy, we inferred the selection coefficient for a range of sample sizes for several different combinations of the population size and selection coefficient. Figure 6 shows a plot of the variance in selection coefficients inferred using Procedures 1 and 3 for sample sizes ranging from *n* = 2 to *n* = 50. For each combination of *N*_*e*_, *s*, and *n*, the trajectories of 100 alleles were simulated under the Wright-Fisher process with an initial allele frequency of *y*_0_ = 0. 1. Samples were taken in generations 50 and 100.

**Figure 6.**
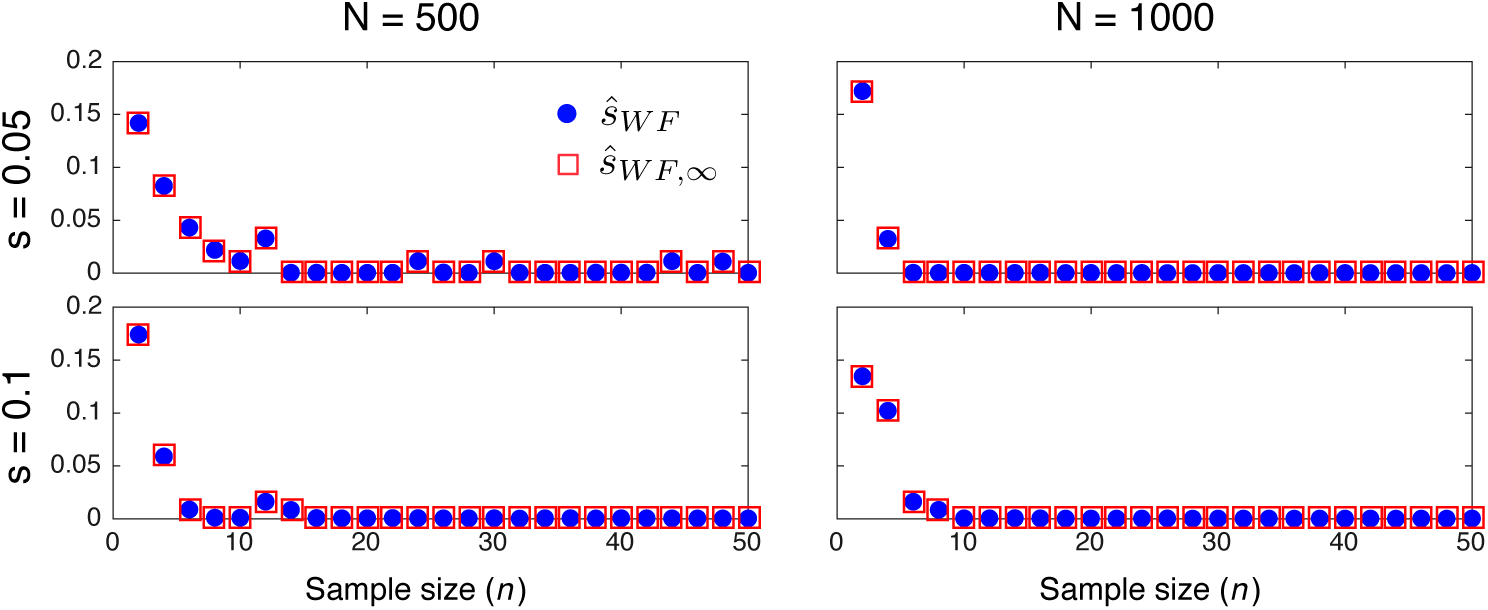
The effect of sample size on inference accuracy. The variance of the estimates produced by the methods in Procedures 1 and 3 are shown for a range of sample sizes.

The plots in Figure 6 suggest that variability due to small sample sizes has a strong effect on the variability in estimates only for sample sizes smaller than 10 alleles. Thus, in all of our simulations we have used a sample size of *n* = 50 alleles so that differences between estimators are not likely to be obscured by the variance in estimates due to small sample sizes.

### 1.9. Computational efficiency of methods.

As we have noted, methods that assume that allele frequency trajectories are deterministic can be considerably faster than methods that account for population size histories. Table 1 shows the average runtimes of the estimators 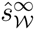, *ŝ*_***W***_, *ŝ*_***D***_, and 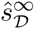 for the computations used to produce Figure 4A-I.

**Table 1.** Mean runtimes of the methods in Figure 4A-I (seconds).

| $N_e$ | $s$ | $\hat{s}_W^\infty$ | $\hat{s}_W$ | $\hat{s}_D^\infty$ | $\hat{s}_D$ |
| --- | --- | --- | --- | --- | --- |
| 100 | 0.01 | 0.01 | 2.30 | 4.74 | 197.25 |
|  | 0.05 | 0.01 | 2.36 | 4.23 | 204.66 |
|  | 0.1 | 0.01 | 2.29 | 3.98 | 217.34 |
| 500 | 0.01 | 0.02 | 134.07 | 4.41 | 185.18 |
|  | 0.05 | 0.01 | 132.45 | 4.41 | 496.83 |
|  | 0.1 | 0.02 | 126.35 | 4.46 | 531.15 |
| 1000 | 0.01 | 0.02 | 175.27 | 4.64 | 196.90 |
|  | 0.05 | 0.02 | 191.53 | 4.78 | 815.27 |
|  | 0.1 | 0.02 | 199.32 | 4.67 | 1950.59 |

From the table, it can be seen that the runtimes are considerably faster for the estimators based on deterministic trajectories (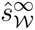 and 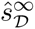). Moreover, the runtimes for 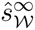 and 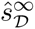 do not depend on the population size or selection coefficient. In comparison, the runtimes for the estimators *ŝ*_***W***_ and *ŝ*_***D***_ increase with increasing *N*_*e*_ and *s* because these methods depend on eigenvalue decompositions or sparse matrix products, which require larger matrices or greater precision when *N*_*e*_ or *s* is large.

## 2. Discussion

The results of our analyses suggest that accurate estimates of selection coefficients from allele frequency time series data can be obtained by assuming that alleles evolve without drift in a population of infinite size. In the majority of our simulations, the estimates obtained using deterministic approximations were nearly as accurate as estimates obtained by explicitly modeling the true population history and they were sometimes more accurate than estimates obtained using crude but reasonable estimates of the population history. Surprisingly, estimates made under the deterministic approximation were generally as accurate as estimates that accounted for drift, due to the fact that the exact maximum likelihood methods had low accuracy when drift was strong.

Accounting for the true population history only resulted in significantly improved estimates of selection coefficients when conditioning on the event that the target allele was segregating in the final sample. Methods that modeled the true population history were more accurate in this case because they could be modified to model conditional trajectories, whereas methods that assumed infinite population sizes could not. These results suggest that methods that account for drift are likely to be preferable under circumstances in which conditioning on segregation is desirable. However, it is important to note that deterministic methods can perform well when population sizes are moderately large if allele frequencies are conditioned on a slightly different event: the event that an allele is found (segregating or fixed) in the final sample.

The idea that ignoring drift can lead to accurate estimates of selection coefficients is not new. In fact, inference methods based on deterministic allele frequency trajectories capitalize on exactly this idea. However, our comparison with estimators based on exact likelihoods makes it possible to characterize the relative loss in accuracy that is incurred when drift is ignored, as well as the demographic, evolutionary, and sampling scenarios under which accounting for drift is likely to be important.

The comparatively accurate estimates achieved by methods that assume deterministic allele frequency trajectories are encouraging for three primary reasons. First, a large number of studies have relied on the assumption that alleles evolve deterministically in order to infer selection coefficients from biological time series data. Our results suggest that these estimates are likely to be nearly as accurate as those obtained using the exact likelihood accounting for drift. Second, estimators based on deterministic trajectories can be considerably faster than estimators that account for drift, making them useful for inferring selection coefficients at large numbers of loci. Third, it may be easier to obtain analytical results under the assumption that allele frequencies change deterministically, simplifying the development of inference methods for inferring selection coefficients under more complicated scenarios; for example, inferring coefficients at linked loci (Illingworth *et al.*, 2012). The ability to model factors such as linkage between alleles under selection may ultimately be more important than modeling drift, as these factors can have a strong effect on evolutionary dynamics (Burke, 2012; Long *et al.*, 2015). Finally, the ability to ignore the population size is useful in situations in which the true population history is unknown or difficult to infer.

In addition to characterizing the degree to which accounting for drift can improve estimates of selection coefficients, our results shed light on the accuracy of exact maximum likelihood methods for inferring selection coefficients from allele frequency trajectories. In accordance with previous work (Schaffer *et al.*, 1977; Gallet *et al.*, 2012), our findings suggest that very small selection coefficients (*s* ≤ 0.01) are difficult to infer if the initial allele frequency and population size are not large. Moreover, even if the population size is large, the accurate inference of a small selection coefficient may require samples taken over hundreds of generations, during which time the selection coefficient could change considerably (Felsenstein, 1976; Siepielski *et al*., 2009).

Despite the difficulties of inferring weak selection coefficients when the population size is small, coefficients of one percent or lower can be inferred accurately if the initial allele frequency is sufficiently high. It is important to note that the selection coefficient need not be high at the time of the very first sampling event, as long as the allele has reached a sufficiently high frequency at one of the intermediate sampling events, leading to quasi-deterministic behavior between some sampling time points that can be exploited by the maximum likelihood estimator. Thus, one need not restrict analyses to cases of selection on standing variation to obtain accurate inferences. Although we have only considered positively selected alleles in our analyses, our results apply equally well to negatively selected alleles, as it is arbitrary whether we choose to track the trajectory of the allele with higher or lower fitness.

We have also considered only low initial allele frequencies (*y*_0_ ≤ 0.1) for selected alleles; however, it is clear from Figure 2 that allele frequency trajectories become increasingly deterministic as the initial allele frequency increases. Thus, the accuracy of a method that assumes a deterministic trajectory will become more similar to that of a method that accounts for drift as the initial allele frequency increases. Conversely, for negatively selected alleles, the accuracy of the deterministic method will approach that of the exact likelihood as the initial allele frequency decreases. Thus, our analyses provide a characterization of inference accuracy for both positively and negatively selected alleles for the full range of starting frequencies.

At first glance, our finding that the population size does not strongly influence estimates of selection coefficients might appear to be at odds with the fact that population size histories can be inferred from allele frequency time series data (O’Hara, 2005; Bollback *et al*., 2008; Ferrer-Admetlla *et al*., 2015). However, this is not the case. Methods for inferring the population size capitalize on information in the short term fluctuations of the allele frequency around its expected value, arising from drift; conversely, estimators of selection coefficients capitalize on the longterm changes in allele frequency due to selection, effectively averaging over the short-term fluctuations due to drift. Our results suggest that allele frequencies often change quasi-deterministically, even in small populations. Thus, deviations around the expected trajectory can be distinguished from long-term changes, allowing effective population sizes to be inferred accurately even in small populations.

We have conducted our analyses under two different models of evolution: the discrete Wright-Fisher model and the continuous diffusion model.

Although the diffusion model was developed as an approximation to the Wright-Fisher process, it also captures the limiting behavior of a large class of evolutionary models, including the Wright-Fisher process, as the population size grows to infinity and mutation and selection parameters are scaled accordingly. Thus, it is reasonable to believe that our findings will generalize to maximum likelihood estimators derived under a wide range of evolutionary models.

Taken together, our results help to characterize the properties of maximum likelihood methods for inferring selection coefficients from time series data. Because of the accuracy and beneficial properties of maximum likelihood methods, it is reasonable to believe that our results provide insight into the accuracy with which it is possible to infer selection coefficients from biological data, and the degree to which accounting for the true population history can improve these estimates. Our results also provide justification for the use of fast inference methods based on the assumption that allele frequencies evolve deterministically. Such methods can be applied to infer selection coefficients efficiently on large genomic data sets with many loci. Finally, our results provide further justification for the use of deterministic approximations in the development of statistical approaches for studying time series data.

## 3. Methods

In this section, we compute the exact probability of an allele frequency trajectory in a population of piecewise-constant size under the discrete Wright-Fisher model and under the diffusion approximation. We also describe how drift can be ignored in these probabilities, yielding approximate estimators of selection coefficients that are similar to commonly-used approaches that assume deterministic allele frequency trajectories.

### 3.1. Computing ℙ_Θ, ***W***_{*O*_[1:*K*]_ = *o*_[1:*K*]_} under the discrete Wright-Fisher model.

To compute the probability ℙ_Θ, ***W***_{*O*_[1:*K*]_ = *o*_[1:*K*]_} under the discrete Wright-Fisher model, we make use of a hidden Markov model (HMM) similar to that presented in Steinrücken *et al*. (2014). However, the hidden state in our discrete model is the count *c*_*t*_ of the number of (unobserved) copies of allele *A* in the population at time *t*, rather than the continuous allele frequency *y*_*t*_.

In our model, the count *c*_*t*_ of allele *A* evolves according to a Wright-Fisher process in which mutation occurs, followed by random mating, selection, and drift. Given that the count of allele *A* in generation *t* is *c*_*t*_ = *i*, let 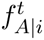 be the frequency of allele A in the gamete pool after mutation. Then

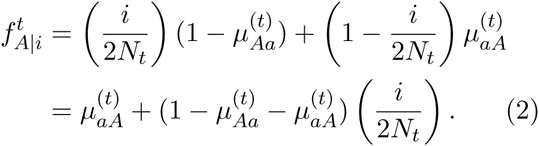

After random mating, the fraction of zygotes with each of the genotypes *AA*, *Aa*, and *aa* is 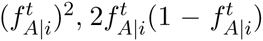, and 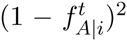, from which it follows that the fraction of genotypes of each kind remaining after selection is given by

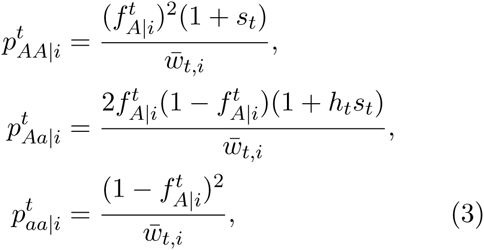

where 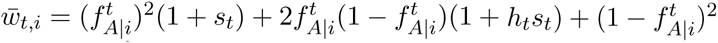 is the mean fitness of the population.

Immediately after selection and before drift occurs, the probability that a randomly chosen allele is of type *A* is given by 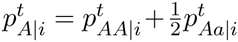.

Then, as the result of drift, the count of allele *A* in generation *t* + 1 is binomially distributed with mean 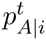. Thus, the probability that allele *A* has count *j* in generation *t* + 1, given that it had count *i* in generation *t* is

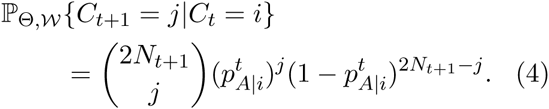

The Wright-Fisher transition matrix **T**_*t*, *t* + 1_ from generation *t* to generation *t* + 1 is the (2*N*_*t*_ + 1) × (2*N*_*t* + 1_ + 1) matrix with entry *i*, *j* given by

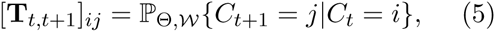

which can be used to obtain the allele frequency distribution at each discrete generation *t* given the initial distribution at some time *s* < *t*. In particular, define *d*_*t*_ = (ℙ{*c*_*t*_ = 0}, ℙ{*c*_*t*_ = 1},…, ℙ{*c*_*t*_= 2*N*_*t*_}), to be the distribution of the count of allele *A* in generation *t*. Using Equation (5), d_*t*_ can be computed recursively as d_*t*_ = 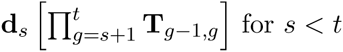.

#### 3.1.1. Computing the probability ℙ_Θ, ***W***_{*O*_[1:*K*]_ = *o*_[1:*K*]_}.

The probability ℙ_Θ, ***W***_{*O*_[1:*K*]_ = *o*_[1:*K*]_} of the observed data is computed using the forward procedure for hidden Markov models. In particular, we define the vector v_*k*_ whose *i*th entry *v*_*k*, *i*_ is the joint probability of the population-wide count of allele *A* at the *k* th sampling event and the observed sample allele counts up to sample *k*:

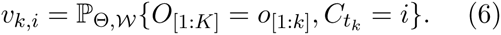

To simplify calculations, we also define the conditional “emission probability”

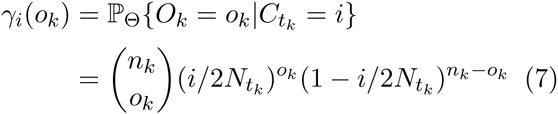

of the observed allele count, conditional on the population allele count. The probability in Equation (7) comes from the fact that the observed allele count at time *t*_*k*_ is a binomial random variable with sample size *n*_*k*_ and probability *c*_*t*_*k*__. We then construct the emission probability vector

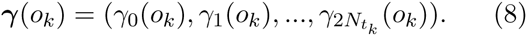

The probability of the data is then given by the forward procedure (Rabiner, 1989), outlined in Procedure 1. In Procedure 1, the formula for v_1_ comes from the fact that

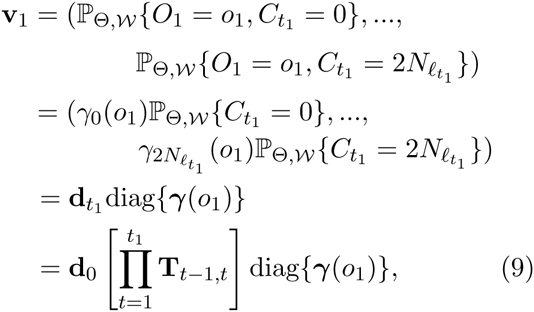

where diag(*γ*) denotes the square matrix whose diagonal entries are given by *γ*.

It has been noted by several authors that computing powers of the transition matrix is computationally prohibitive, providing one motivating factor for the use of approximations of the Wright-Fisher process, such as the diffusion and Gaussian approximations (Ewens, 1963; Feder *et al*., 2014; Lacerda and Seoighe, 2014). However, the products 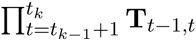 in Procedure 1 do not require products of the transition matrix **T**_*t*–1, *t*_,because it suffices to repeatedly compute vector-matrix products instead of multiplying full matrices together. In practice, this can be done very quickly, even for large population sizes. A similar fast procedure was carried out by Zhao *et al*. (2014) to simulate trajectories under the Wright-Fisher model.

### 3.2. Computing ℙ_Θ, ***D***_{*O*_[1:*K*]_ = *o*_[1:*K*]_} under the diffusion approximation.

The diffusion approximation models the evolution of the continuous population frequency *Y*_*t*_ of allele *A*, rather than its count *C*_*t*_. The time-evolution of the random frequency *Y*_*t*_ is governed by the diffusion transition density *p*_Θ_(*s*, *t*; *x*, *y*) given by

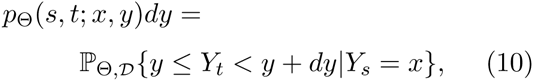

for an infinitesimal increment *dy*. The quantity *p*_Θ_(*s*, *t*; *x*, *y*) specifies the density of the allele frequency at time *t*, conditional on the value of the allele frequency at an earlier time *s*. For more details about the transition density function of the diffusion approximation, see Appendix A.

Using the diffusion transition density *p*_Θ_(*s*, *t*; *x*, *y*) Steinrücken *et al*. (2014) developed an HMM to compute the probability ℙ_Θ, ***D***_{*O*_[1:*K*]_ = *o*_[1:*K*]_} of the data in a single epoch of constant size by efficiently integrating over the hidden allele frequencies {*yt*_1_,…, *yt*_*K*_} at the set of sampling times. Here, we extend this HMM to the case of piecewise-constant population size.

To compute the probability ℙ_Θ, ***D***_{*O*_[1:*K*]_ = *o*_[1:*K*]_} efficiently, Steinrücken *et al*. (2014) define the quantities *f*_*k*_(*y*) and *g*_*k*_(*y*) satisfying

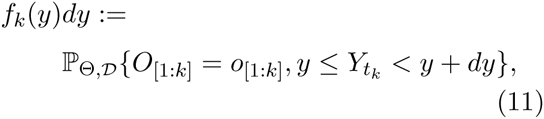

and

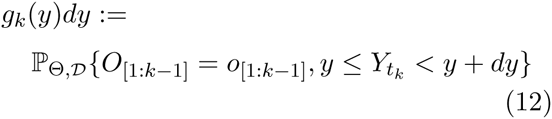

for an infinitesimal increment *dy*. The quantity *f*_*k*_(*y*) is the joint density of the allele frequency at time *t*_*k*_ and the observed counts up to sampling event *k*. The quantity *g*_*k*_(*y*) is the joint density of the allele frequency at time *t*_*k*_ and the observed counts up to sampling event *k* – 1.

It follows from the definition of *f*_*k*_(*y*) that the probability of the data is given by

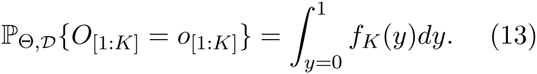

The quantity *f*_*K*_(*y*) can be obtained efficiently by recursion using the relationships

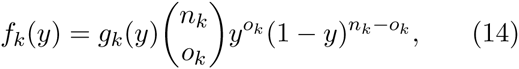

and

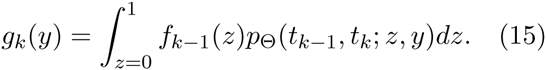

Equation (14) follows from the fact that the observed number of copies of allele *A* at sampling event *k* is binomially distributed with count *n*_*k*_ and probability *y*_*t*_*k*__ and Equation (15) follows from the law of total probability integrating over *Y*_*t*_*k*__ 1.

Let *B*_*ℓ*, *i*_(*y*) (*i* = 0,1,…) be the *i*th eigenfunction of the backward diffusion operator ***𝓛***_*ℓ*_ and let *π*_*ℓ*_(*y*) be the speed density of ***𝓛***_*ℓ*_ (Appendix A). Steinrücken *et al*. (2014) demonstrated that the recursive formulas in Equations (14) and (15) can be evaluated efficiently by expressing *f*_*k*_(*y*) and *g*_*k*_(*y*) as series of the form

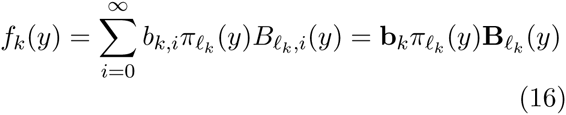

and

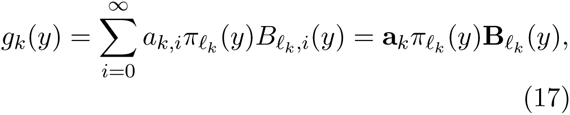

where **B**_*ℓ*_(*y*) = (*B*_*ℓ*, 0_(*y*), *B*_*ℓ*, 1_(*y*),…) and where b_*k*_ = (*b*_*k*, 0_, *b*_*k*, 1_,…) and a_*k*_ = (*a*_*k*, 0_, *a*_*k*, 1_,…) are vectors of constants that encode the densities *f*_*k*_(*y*) and *g*_*k*_(*y*) at the beginning of the epoch. In Appendix B, we extend the results of Steinrücken *et al*. (2014) to derive recursive formulas for the coefficients *a*_*k*_ and *b*_*k*_, resulting in Procedure 2, which computes the probability of an allele frequency trajectory under the diffusion approximation in a population of piecewise constant size.

### 3.3. Conditional probabilities.

Sometimes it is desirable to compute the probability of the observed allele counts conditional on the event *S*_*K*_ that allele *A* is segregating in the final sample. In this section, we provide formulas for these conditional probabilities under the Wright-Fisher and diffusion models.

#### 3.3.1. Computing ℙ_Θ, ***W***_{*O*_[1:*K*]_ = *o*_[1:*K*]_}.

In Section 1.7.1, we consider the probability ℙ_Θ, ***W***_{*O*_[1:*K*]_ = *o*_[1:*K*]_} of the data conditional on the event *S*_*K*_ that allele *A* is segregating in the final sample. In Appendix C, we show that in the case of the discrete Wright-Fisher model,

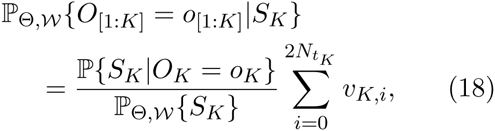

where *v*_*k*, *i*_ is defined in Equation (6) and ℙ{*S*_*K*_|*O*_*K*_ = *o*_*K*_} = 1 if 1 ≤ *o*_*K*_ < *n*_*K*_, or 0 otherwise. The probability ℙ_Θ, ***W***_{*S*_*K*_} is given in Equation (C.3). Thus, if we wish to compute conditional probabilities under the Wright-Fisher model, we carry out Procedure 1, replacing step 3 with Equation (18).

#### 3.3.2. Computing ℙ_Θ, ***W***_{*O*_[1:*K*]_ = *o*_[1:*K*]_}|*F*_*K*_}.

Similarly, for the event *F*_*K*_ that allele *A* is segregating or fixed in the final sample, we show in Appendix C that

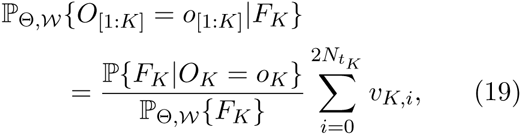

where *v*_*k*, *i*_ is defined in Equation (6) and ℙ{*F*_*K*_|*O*_*K*_ = *o*_*K*_} = 1 if 1 ≤ *o*_*K*_ ≤ *n*_*K*_, or 0 otherwise. The probability ℙ_Θ, ***W***_{*F*_*K*_} is given in Equation (C.6). If we wish to compute conditional probabilities under the Wright-Fisher model, we carry out Procedure 1, replacing step 3 with Equation (19).

#### 3.3.3. Computing ℙ_Θ, ***D***_{*O*_[1:*K*]_ = *o*_[1:*K*]_|*S*_*K*_}.

In the case of the diffusion approximation, we show in Appendix D that the conditional probability of the data given *S*_*K*_ can be computed as

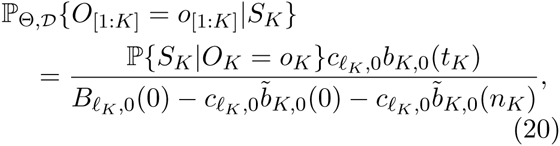

where ℙ{*S*_*K*_|*O*_*K*_ = *o*_*K*_} = 1 if 1 ≤ *o*_*K*_ ≤ *n*_*K*_ or 0 otherwise, and

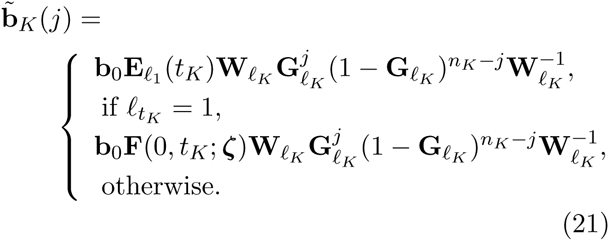

Thus, if we are interested in conditional probabilities under the diffusion model, we carry out Procedure 2, replacing step 3 with Equation (20).

### 3.4 The probability in the absence of genetic drift.

If we ignore genetic drift, the allele frequency changes deterministically over time. Here, we obtain versions of Procedures 1 and 2 in the case when the changes in allele frequency arising from genetic drift are negligible relative to the changes due to selection and recurrent mutation.

#### 3.4.1. Deterministic allele frequency trajectories under the Wright-Fisher model.

If there is no contribution to the change in allele frequency arising from genetic drift, the allele frequency in a given generation is equal to its expectation after mutation, random mating, and selection, conditional on its value in the previous generation. Because the expectation is not necessarily integer-valued, we no longer consider discrete integer allele counts *c*_*t*_. Instead, we track the expected allele frequency, which we denote by 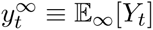, where the subscript ∞ denotes the expectation when drift is negligible.

The expected frequency 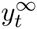 is obtained by combining Equations (2) and (3), ignoring the drift step in Equation (4), yielding

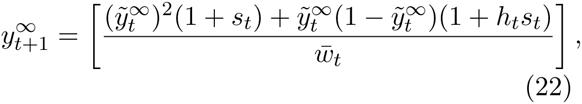

where

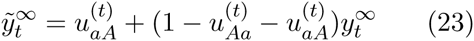

and 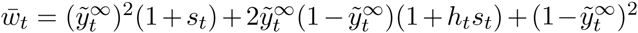 Equations (22) and (23) are iterated to find the allele frequency in any generation *t* > 0.

#### 3.4.2. Deterministic allele frequency trajectories under the diffusion model.

Under the diffusion model in an Epoch *ℓ* of constant size, the allele frequency *Y*_*t*_ obeys the stochastic differential equation (SDE)

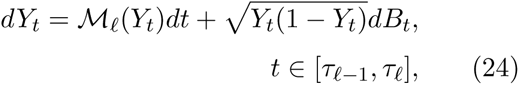

with the initial condition *Y*_*τ*_*ℓ*–1__ = *y*_*τ*_*ℓ*–1__, where time is measured in units of generations and *τ*_*ℓ*–1_ is the time at which Epoch *ℓ* begins (Durrett, 2008, Section 7.2). The quantity 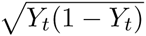 in Equation (24) controls random fluctuations due to drift whereas the quantity ***M***_*ℓ*_(*y*) describes the deterministic change in the mean frequency of the allele over time due to mutation and selection and is given by

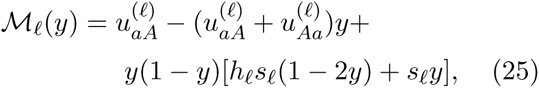

In Equation (25) we have rescaled the usual form of ***M***_*ℓ*_ so that time is measured continuously in units of generations.

If the drift term in Equation (24) is negligible compared with ***M***_*ℓ*_(*Y*_*t*_), then Equation (24) can be approximated by the ordinary differential equation

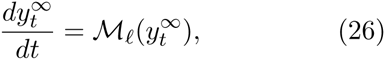

where we may write 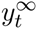 instead of *Y*_*t*_ because the evolution of the allele frequency is deterministic and follows its expectation in the absence of drift.

We can also suppress the explicit dependence on the epoch *ℓ* by defining 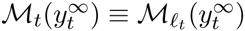, yielding

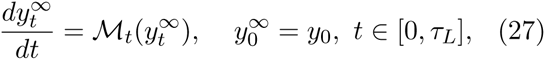

which holds for the full population history across all epochs *ℓ* = 1,…, *L*. Equation (27) can be solved numerically, for instance by choosing a sufficiently small time step Δ*t* and iteratively computing 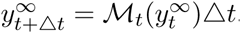.

#### 3.4.3. Sample probabilities based on deterministic allele frequency trajectories.

To compute the probability 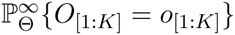 under either the discrete Wright-Fisher or diffusion models when drift is negligible, we note that the observations (*O*_1_,…, *O*_*K*_) are conditionally independent of one another, given the underlying allele frequencies.

Thus, in the absence of drift we have

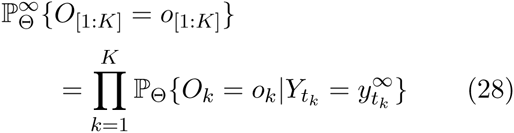

for both the diffusion and Wright-Fisher models, where 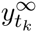 is the deterministic allele frequency at time *t*_*k*_, for *k* = 1,…, *K*. Using Equations (22) and (28), the probability of the data under the Wright-Fisher model in a population without drift can be obtained using Procedure 3. Similarly, using Equations (27) and (28), the probability of the data in the case of the diffusion model is given by Procedure 4.

## Acknowledgments

This research was supported by the National Institutes of Health (grant number R01-GM094402) and by a Packard Fellowship for Science and Engineering.

## Appendix A.

Diffusion transition densities: background

The equations in Section 3.2 were derived under a model in which the selected allele A evolves under the diffusion approximation in a population of piecewise constant size. Given that allele *A* has frequency *x* at a fixed time *s*, the density at a later time *t* is given by the transition density of the diffusion approximation (Equation 10). Steinrücken *et al*. (2014) derived a formula for the density for the case of a single population of constant size. Here, we review this derivation to provide background and notation for the derivation of the diffusion model probability computed in Procedure 2.

### A.1 The diffusion approximation in a population of constant size.

Let *p*_*ℓ*_(*s*, *t*; *x*, *y*) denote the transition density restricted to a specific epoch *ℓ* of constant size with *s*, *t* ∈ *ℓ*. The density *p*_*ℓ*_(*s*, *t*; *x*, *y*) is the unique solution of the Kolmogorov backward equation,

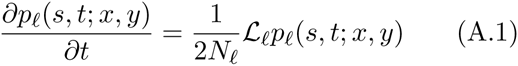

satisfying the terminal condition *ρ*_*s*_(*y*) = *δ*(*y* ‒ *x*), where *δ*(·) is the Dirac delta distribution and ***𝓛***_*ℓ*_ is the Kolmogorov backward operator in the epoch defined in Equation (A.2). The factor 1/2N_*ℓ*_ in Equation (A.1), is introduced so that the timescaling is the same in all epochs, and time is measured continuously in units of generations.

The Kolmogorov backward operator is defined in terms of the scaled mutation and selection parameters 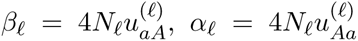, and *σ*_*ℓ*_ = *N*_*ℓ*_*s*_*ℓ*_ as

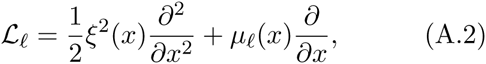

where the quantity

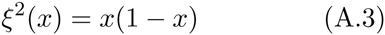

captures the contribution to the change in allele frequency arising from genetic drift and

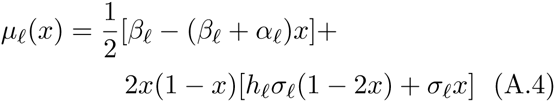

captures the contribution from recurrent mutation and selection.

Song and Steinrücken (2012) showed that *p*_*ℓ*_(*s*, *t*; *x*, *y*) can be expressed as an expansion in the eigenfunctions of ***𝓛*** of the form

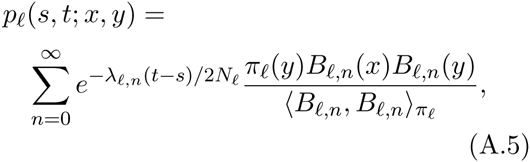

where 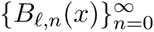 are the eigenfunctions of ***𝓛***_*ℓ*_ with associated eigenvalues 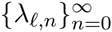 and the function *π*_*ℓ*_(*y*) is given by

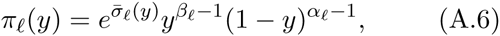

where *σ*̄_*ℓ*_(*y*) = 4*h*_*ℓ*_*σ*_*ℓ*_*y*(1 – *y*) + 2*σ*_*ℓ*_*y*^2^. The inner product 〈*f*, *g*〉_*ω*_ with respect to a weight function *ω*(*x*) in Equation (A.5) is defined for two functions *f* and *g* on an interval [*a*, *b*] by

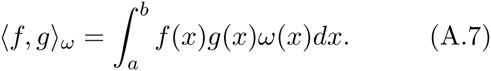

In Equation (A.5), the inner product 〈·, ·〉_*π*_*ℓ*__ is taken over the interval [0,1] with respect to *π*_*ℓ*_(*y*).

### A.2. Expressions for the quantities in Equation (A.5).

Expressions for the eigenvalues 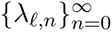, eigenfunctions 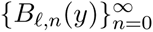, and inner products 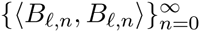 in Equation (A.5) can be obtained using a matrix formulation developed by Steinrücken *et al*. (2014). In particular, the eigenfunctions 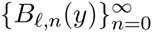 can be expressed as

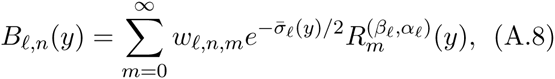

where 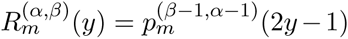 and 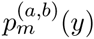 is the *m*th classical Jacobi polynomial (Abramowitz and Stegun, 1972, Chapter 22). The vector *w*_*ℓ*, *n*_ = (*w*_*ℓ*, *n*, 0_, *w*_*ℓ*, *n*, 1_,…) is the *n*th left eigenvector of the infinite-dimensional matrix

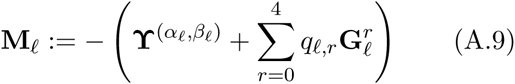

corresponding to the *n*th eigenvalue *λ*_*ℓ*, *n*_, where 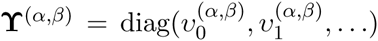 is the diagonal matrix with elements given by 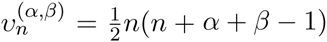 and the quantities *q*_*ℓ*, *r*_ are given by

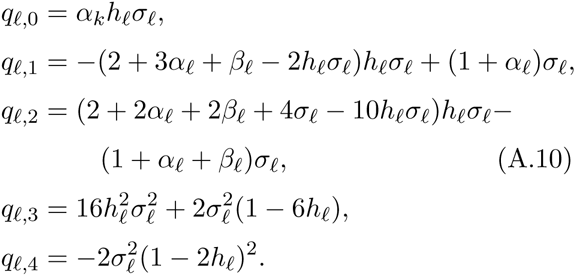

The matrix 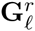 in Equation (A.9) has elements given by

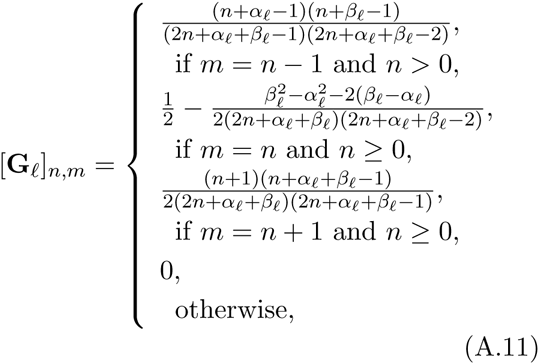

which correspond to the coefficients of the three-term recurrence relation satisfied by the Jacobi Polynomials.

### A.3. Matrix expressions for the transition density.

It is computationally and notationally simpler to express the eigenfunctions of ***𝓛***_*ℓ*_ and the transition density as products of matrices. In particular, we can express Equation (A.8) as

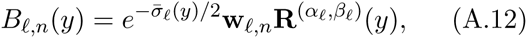

where

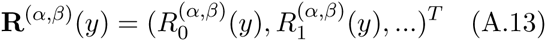

and we can express the vector **B**_*ℓ*_(*y*) of eigenfunctions as

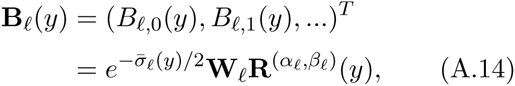

where

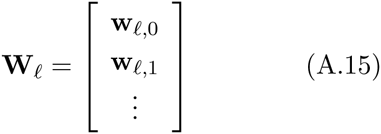

is the matrix whose rows are the left eigenvectors of the matrix **M**_*ℓ*_ in Equation (A.9).

Using Equations (A.5) and (A.14), the transition density in a single epoch *ℓ* can then be expressed as the matrix product

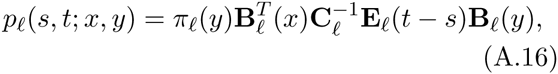

where

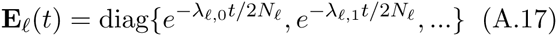

and 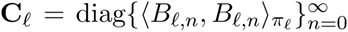. Steinrücken *et al*. (2014) showed that the matrix **C**_*ℓ*_ in Equation (A.16) can be expressed as

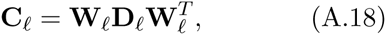

where

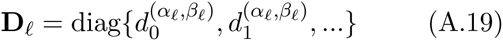

and

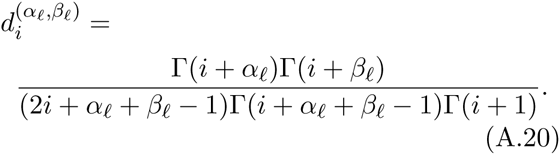

Thus, the transition density in a single epoch can be computed by constructing matrix **M**_*ℓ*_, computing its eigenvectors **W**_*ℓ*_ and eigenvalues (*λ*_*ℓ*, 0_, *λ*_*ℓ*, 1_,…), and plugging these into the components of Equation (A.16). In practice, because the matrix **M**_*ℓ*_ has infinite dimension, we approximate it by truncating its dimensions at some large integer *M* yielding approximate eigenvectors 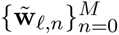 and eigenvalues 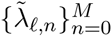. We also truncate the length of the vector **B**_*ℓ*_(*y*) at a large integer *N*. Although these truncations lead to approximate values of the transition density, the approximation can be made arbitrarily precise by taking *N* ≤ *M* to be sufficiently large.

## Appendix B. Recursions for the coefficients *a*_*k*_ and *b*_*k*_.

### B.1. Discussion of the problem.

Here, we extend the HMM of Steinrücken *et al*. (2014) to accommodate populations of piecewise constant size. As we noted in Section 3.2, the probability ℙ_Θ, ***D***_{*O*_[1:*K*]_ = *o*_[1:*K*]_} of the data under the diffusion model can be obtained using the equation

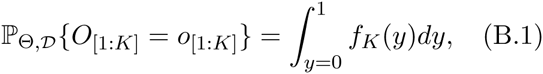

where the quantity *f*_*K*_(*y*) is obtained by recursively evaluating Equations (14) and (15). Because *f*_*k*_(*y*) and *g*_*k*_(*y*) can be expressed as the series *f*_*k*_(*y*) = *π*_*ℓ*_*k*__(*y*)b_*k*_B_*ℓ*_*k*__(*y*) and *g*_*k*_(*y*) = *π*_*ℓ*_*k*__(*y*)a_*k*_B_*ℓ*_*k*__(*y*) (Equations 16 and 17), determining *f*_*k*_(*y*) and *g*_*k*_(*y*) amounts to determining the coefficients *a*_*k*_ and *b*_*k*_. Thus, it is useful to develop analogs of the recursions (13) and (14) that apply to the coefficients themselves.

### B.2. Equations for propagating coefficients.

From Equation (14), it can be seen that obtaining *f*_*k*_(*y*) from *g*_*k*_(*y*) involves only multiplication by a polynomial in *y*. Thus, the formula for obtaining the coefficients *b*_*k*_ from the coefficients *a*_*k*_ does not depend on the population history and, therefore, it can be obtained from results in Steinrücken *et al*. (2014) who derived formulas for the recursion for the case of a population of constant size. However, the formula for obtaining *g*_*k*_(*y*) from *f*_*k*–1_(*y*) (Equation 15) involves the transition probability *p*_Θ_(*t*_*k*–1_, *t*_*k*_; *z*, *y*), which depends on the population parameters Θ. Thus, it is necessary to account for the population history when computing the coefficients *a*_*k*_ from the coefficients *b*_*k*–1_.

To obtain *a*_*k*_ from *b*_*k*–1_, we first consider the more general problem of obtaining the generalized vector of coefficients *a*_*k*_(*t*) from *b*_*k*–1_, where *a*_*k*_(*t*) is defined as the vector of coefficients of the expansion of the generalized density *g*_*k*_(*y*, *t*) defined by

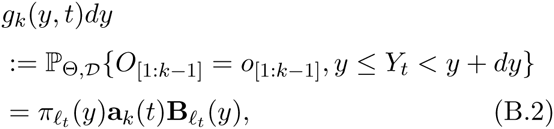

i.e., the joint density of the observed data up to sample *k* – 1 and the allele frequency at time *t*, where we assume *t*_*k*–1_ ≤ *t* so that the time *t* at which *g*_*k*_(*y*, *t*) is evaluated occurs later than the time *t*_*k*–1_ at which *f*_*k*_(*y*) is evaluated. The generalized density *g*_*k*_(*y*, *t*) is related to the density *g*_*k*_(*y*) defined in Equation (12) by *g*_*k*_(*y*) = *g*_*k*_(*y*, *t*_*k*_).

To obtain *a*_*k*_(*t*) from *b*_*k*–1_, there are two scenarios to consider: the case in which both *t*_*k*–1_ and *t* lie within the same epoch *ℓ* and the case in which *t*_*k*–1_ and *t* lie within distinct epochs. Our derivations of these separate cases provide the results necessary for step 2 of Procedure 2.

#### B.2.1. The case *ℓ*_*t*_*k*–1__ = *ℓ*_*t*_ = *ℓ*.

If both *t*_*k*–1_ and *t* lie within the same epoch *ℓ*, then the transition density is given by Equation (A.16) and we have

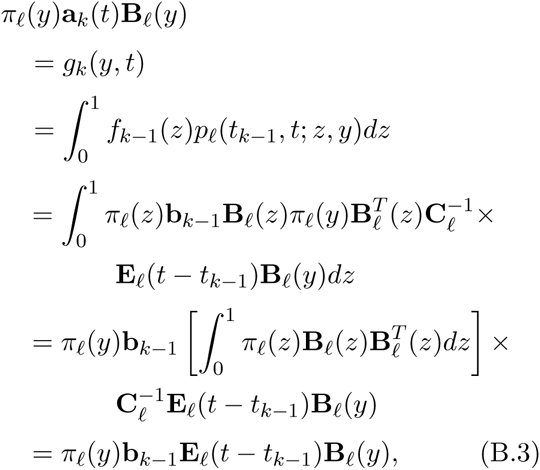

where the second equality follows from Equation (15) and where we have used the fact that 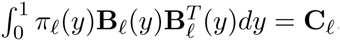. Because the eigenfunctions 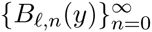 form a complete basis of the Hilbert space defined with respect to the inner product 〈·, ·〉_*π*_*ℓ*__, the coefficients in the expansion on the left-hand side of Equation (B.3) must equal those on the right-hand side. Thus,

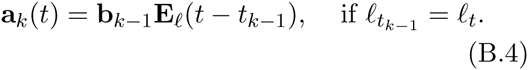

#### B.2.2. The case when *ℓ*_*t*_*k*–1__ ≠ *ℓ*_*t*_.

If the times *t*_*k*–1_ and *t* lie in different epochs, *∓*_*t*_*k*–1__ and *ℓ*_*t*_, then the transition density is no longer given by Equation (A.16). Instead, we must use a formula for the transition density across multiple epochs of different sizes. Steinrücken *et al*. (2015) showed that if the allele frequency density *ρ*_*ℓ*, *s*_(*y*) at time s in epoch *ℓ* is given by the expansion

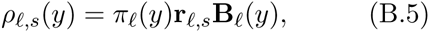

where r_*ℓ*, *s*_ = (*r*_*ℓ*, *s*, 0_, *r*_*ℓ*, *s*, 1_,…) are the coefficients encoding the density at time *s* in the basis of the eigenfunctions 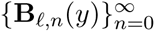, then at time t in epoch *ℓ* + 1, the allele frequency density is given by *ρ*_*ℓ* + 1, *t*_(*y*) = *π*_*ℓ* + 1_(*y*)r_*ℓ* + 1, *t*B_*ℓ* + 1_(*y*), where the coefficients r_*ℓ*_ + 1, *t*_ are given by

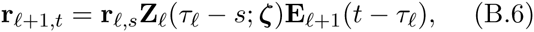

where *τ*_*ℓ*_ is the time of the terminating boundary of epoch *ℓ* and

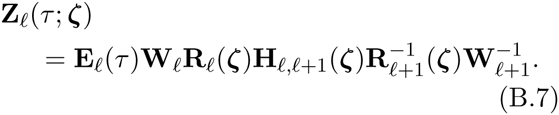

In Equation (B.7), **R**_*ℓ*_(*ζ*) and **H**_*ℓ*, *ℓ* + 1_(*ζ*) are given by

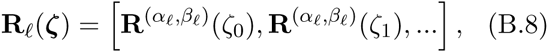

where **R**^*α*, *β*^(*y*) is defined in Equation (A.13) and

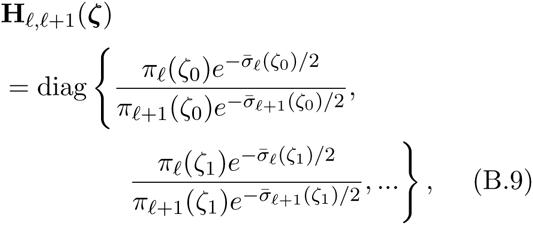

for an arbitrary collection of distinct values *ζ* = (*ζ*_0_, *ζ*_1_,…) ∈ [0,1]. In practice, we take *ζ* to be the Chebyshev nodes (Steinrücken *et al*., 2015).

By repeated application of Equation (B.6), it follows that if the coefficients **r**_ℓ_*s*_, *s*_ encode the density *ρ*_*s*_(*y*) at time *s* in epoch *ℓ*_*s*_, then the coefficients **r**_*ℓ*_*t*_, *t*_ encoding the density *ρ*_*t*_(*y*) at time *t* in epoch *ℓ*_*t*_ > *ℓ*_*s*_ are given by *r*_*ℓ*_*t*_, *t*_ = **r**_*ℓ*_*s*_, *s*_**F**(*s*, *t*; *ζ*), where

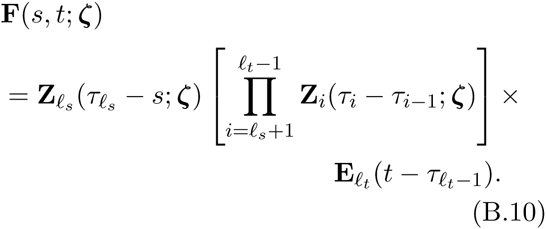

Moreover, if we define **r**_*ℓ*_*s*_, *s*_(*x*) to be the vector of coefficients encoding the density *ρ*(*y*) = *δ*(*y* – *x*), then it follows from Equation (B.10) that the transition density *p*_Θ_(*s*, *t*; *x*, *y*) for times *s* < *t* lying in distinct epochs *ℓ*_*s*_ < *ℓ*_*t*_ is given by

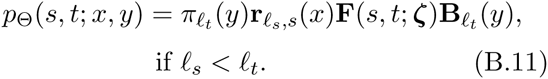

For the initial condition *ρ*_*ℓ*, *s*_(*y*) = *δ*(*y* – *x*), it was shown in Proposition 1 of Steinrücken *et al*. (2014) that the coefficients **r**_*ℓ*_*s*_, *s*_(*x*) are given by

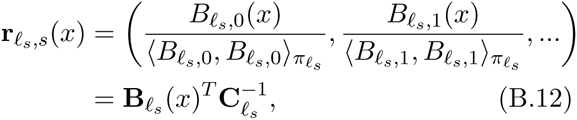

yielding

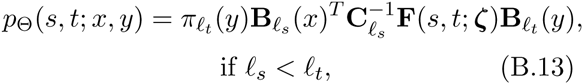

which is obtained by plugging Equation (B.12) into Equation (B.11).

We can now plug Equation (B.13) into Equation (15) to obtain a relationship between a_*k*_(*t*) and *b*_*k* – 1_ when times *t*_*k* – 1_ and *t* lie in different epochs:

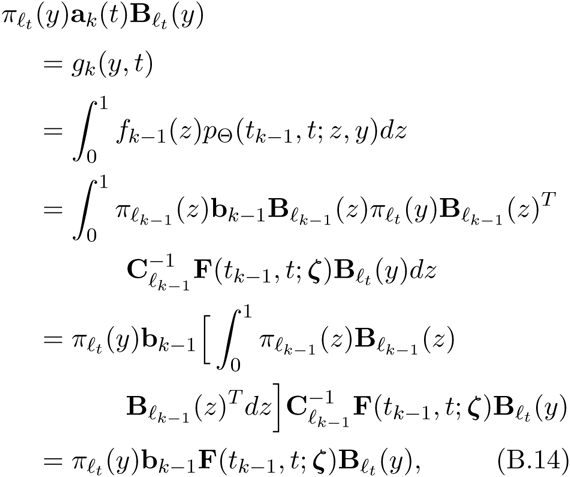

where we have again used the fact that 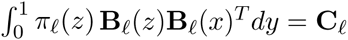. Finally, by the uniqueness of expansions in the Hilbert basis 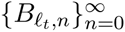, we have

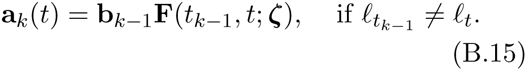

The results derived in Section B.2.2 provide the machinery necessary to propagate the coeffcients ak and bk in the HMM over time. These results can now be used to compute the probability of observing a set of sampled allele frequencies under the diffusion model.

### B.3. Derivation of lemmas necessary for Procedure 2.

We now obtain three lemmas that provide the steps in Procedure 2.

#### Lemma B.3.1.

If the initial frequency density *ρ*_0_(*y*) at time *t*_0_ = 0 is *ρ*_0_(*y*) = δ(*y* – *x*), then the value of the initial vector b_0_ encoding the quantity *f*_0_(*y*) is given by

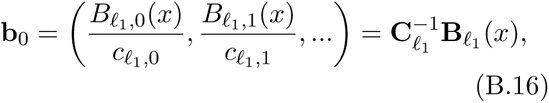

where **B**_ℓ_(*x*) is given in Equation (A.14) and **C**_*ℓ*_ is the diagonal matrix given in Equation (A.18).

*Proof*. Because b_0_ depends only on the parameters Θ_*ℓ*_1__ in the first epoch, the proof of Lemma B.3.1 is the same whether we consider a population composed of a single epoch, or a population composed of multiple epochs. The equation for *f*_*k*_(*y*) (Equation 16) is the same as Equation 2.14 of Steinrücken *et al*. (2014). Thus, the coefficients b_*k*_ in this paper correspond to the coefficients b_*k*_ in Steinrücken *et al*. (2014) who proved Lemma B.3.1 for the case of a population of constant size.

Thus, the first equality in Lemma B.3.1 follows directly from Proposition 1 of Steinrücken *et al*. (2014). The matrix representation in the second equality follows directly from the definitions of **C**_*ℓ*_ and **B**_*ℓ*_(*x*).

#### LemmaB.3.2.

Let **G**_*ℓ*_, **W**_*ℓ*_, **E**_*ℓ*_(*t*), and **F**(*s*, *t*; *ζ*) denote the matrices defined in Equations (A.11), (A.15), (A.17), and (B.10), respectively, where **ζ** = (ζ_0_, ζ_1_,…) *is a set of distinct values arbitrarily chosen such that* {ζ_0_, ζ_1_,…} ∈ [0,1]. Then the coefficient vectors **a**_*k*_ and **b**_*k*_ satisfy the recursive relationships

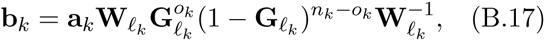

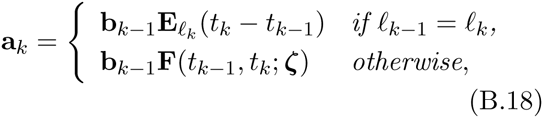

where 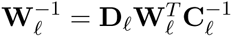

*Proof*. The relationship in Equation (B.18) is obtained immediately by setting *t* = *t*_*k*_ in Equations (B.4) and (B.15), which follows because a_*k*_(*t*_*k*_) = a_*k*_. The relationship in Equation (B.17) does not depend on the population parameters Θ; therefore, Equation (B.17) is the same as that derived in Steinrücken *et al*. (2014), who considered a population of constant size (see Steinrücken *et al*. (2014), Theorem 2).

#### Lemma B.3.3.

The probability ℙ_Θ, ***D***_{*O*_[1:*K*]_ = *o*_[1:*K*]_} of observing the allele counts *o*_[1:*K*]_, given the population parameters Θ is

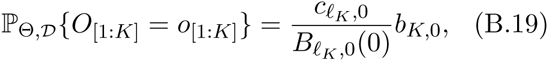

where *c*_*ℓ*, 0_ = [**C**_*ℓ*_]_0, 0_ is element 0, 0 of the matrix **C**_*ℓ*_ in Equation (A.18) and

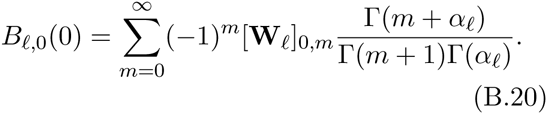

The quantity [**W**_*ℓ*_]_*i*, *j*_ in Equation (B.20) is element *i*, *j* of the matrix **W**_*ℓ*_ given in Equation (A.15).

*Proof*. Equation (B.19) can be obtained by integrating over the joint density *f*_*K*_(*y*) of the data *O*_[1:*K*]_ and the allele frequency *Y*_*t*_*K*__ at the final sampling time:

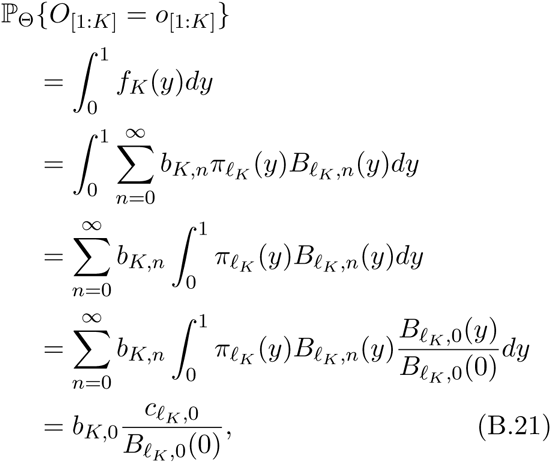

where *c*_*ℓ*_*k*_, 0_ = [C_*ℓ*_*K*__]_0, 0_ ≡ 〈*B*_*ℓ*_*K*_, 0_, *B*_*ℓ*_*K*_, 0_〉_*π*_*ℓ*_*K*___. In the fourth equality we have used the fact that *B*_*ℓ*, 0_(*y*) = *B*_*ℓ*, 0_(0) is a constant function in *y*. To see why *B*_*ℓ*, 0_(*y*) is constant, note that the eigenvalues *λ*_*ℓ*, 0_, *λ*_*ℓ*, 1_,…are non-negative and strictly increasing. Thus, all terms in Equation (A.5) must vanish in the limit *s* → –∞, except possibly the term *n* = 0. Because *p*_*ℓ*_(*s*, *t*; *x*, *y*) approaches the stationary density in the limit *s* → –∞, it must be the case that *λ*_*ℓ*, 0_ = 0, so at least one term does not vanish. Thus, we have

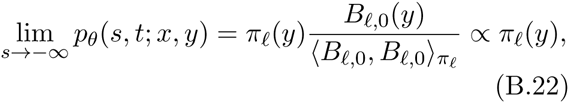

where we have used the fact that *π*_*ℓ*_(*y*) is proportional to the stationary density of the diffusion equation in Epoch *ℓ* It follows from Equation (B.22) that *B*_*ℓ*, 0_(*y*) is constant. Thus, we obtain the result, proving Equation (B.19). Equation (B.20) follows directly from the proof of Proposition 3 in Steinrücken *et al*. (2014).

## Appendix C. Conditional probabilities: the Wright-Fisher model

Under the Wright-Fisher model, the probability ℙ_Θ, ***W***_{*O*_[1:*K*]_ = *o*_[1:*K*]_|*S*_*K*_} of the observed allele counts, conditional on the event *S*_*K*_ that allele *A* is segregating in the final sample can be computed using the fact that

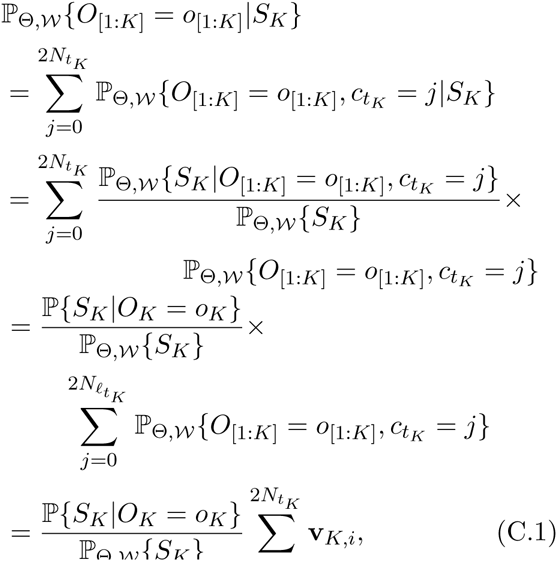

where the third equality in Equation (C.1) follows from the fact that the conditional probability ℙ_Θ, ***D***_{*S*_*K*_|*O*_[1:*K*]_ = *o*_[1:*K*]_, *c*_*t*_*K*__ = *j*} depends only on the allele count *o*_*K*_ and the final equality in Equation (C.1) follows from the definition of **v**_*k*_.

The probability ℙ{*S*_*K*_|*O*_*K*_ = *o*_*K*_} in Equation (C.1) is given by

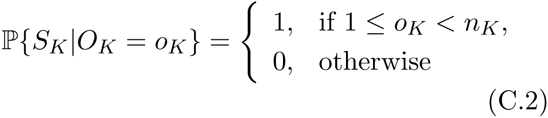

and the probability ℙ_Θ, ***W***_{*S*_*K*_} is given by

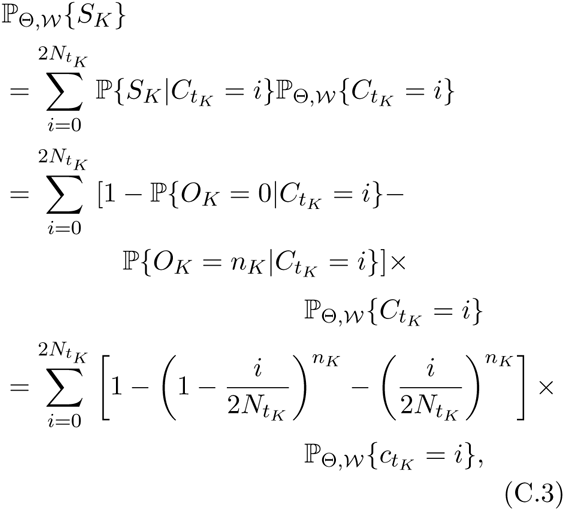

where, as before, ℙ_Θ, ***W***_{*C*_*t*_*K*__ = *i*} is given by the *i*th element of 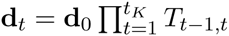.

Note that it is easy to condition on other configurations of the final sample using a procedure similar to that used to derive Equation (C.1). For example, for the event FK that allele *A* is segregating or fixed in the final sample, which we consider in Section 1.7.2, the probability ℙ_Θ, ***W***_{*O*_[1:*K*]_ = *o*_[1:*K*]_|*F*_*K*_} is given by

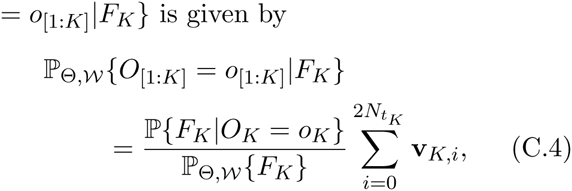

where

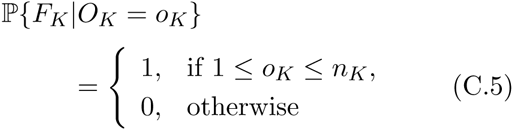

and

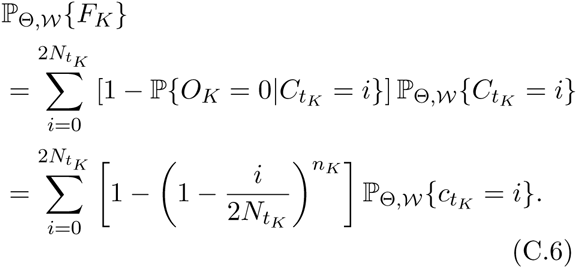

Other probabilities can be obtained in a similar fashion.

## Appendix D. Conditional probabilities: diffusion model

Under the diffusion approximation, the probability ℙ_Θ, ***D***_{*O*_[1:*K*]_ = *o*_[1:*K*]_|*S*_*K*_} of the observed allele counts conditional on the event *S*_*K*_ that allele *A* is segregating in the final sample can be computed using the fact that

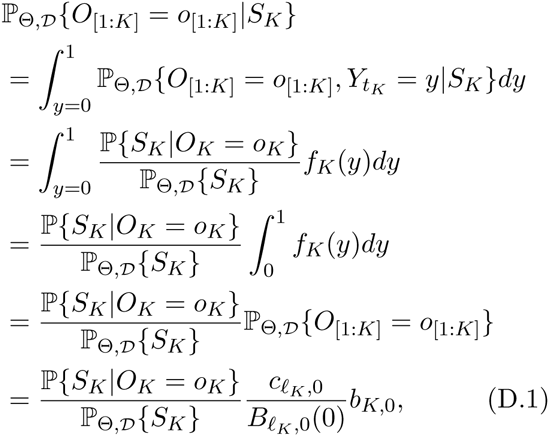

where the second equality follows from the fact that the conditional probability ℙ{*S*_*K*_|*O*_*K*_ = *o*_*K*_, *Y*_*t*_*K*__ = *y*} depends only on the allele count *o*_*K*_ in the final sample, and the final equality follows from Equation (B.19).

The probability ℙ_Θ, ***D***_{*S*_*K*_} can be computed as

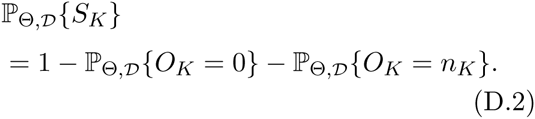

In Equation (D.2), the probability ℙ_Θ, ***D***_{*O*_*K*_ = *j*} can be found easily by noting that if the only sampling time is *t*_*K*_, at which *O*_*K*_ = *j* lineages are observed, then the probability computed using Procedure 2 is precisely the probability ℙ_Θ, ***D***_{*O*_*K*_ = *j*}.

Consider the problem in which the only sampling occurs at time *t*_*K*_ and denote the coefficient vectors for this related problem by **a**̃_*k*_ and **b**̃_*k*_. Then, by Equation (B.19), we see that

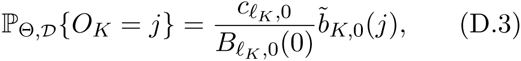

where *b*̃_*K*_(*j*) is is obtained by computing the steps in Procedure 2. In Step 1, we compute

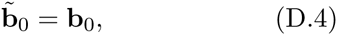

which follows because the initial vector **b**_0_ depends only on the initial frequency. In Step 2, we compute

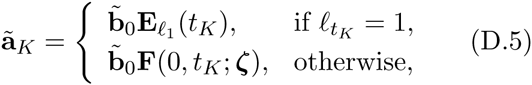

which follows because the coefficients are propagated directly from time *t*_0_ = 0 to time *t*_*K*_. Finally, in Step 3 we have

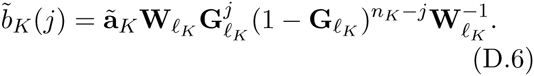

Combined together, Equations (D.4), (D.5), and (D.6) yield

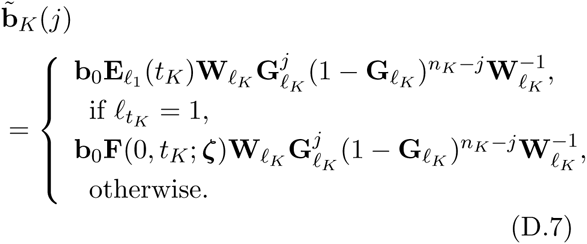

Plugging Equations (D.2) and (D.3) into Equation (D.1) gives

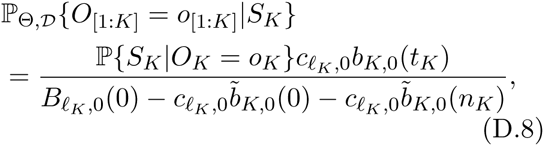

where

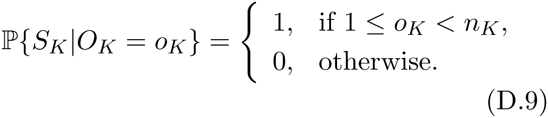

Note that it is easy to condition on other configurations of the final sample by computing the probabilities ℙ{*V*_*K*_|*O*_*K*_ = *o*_*k*_} and ℙ_Θ, ***W***_{*V*_*K*_} for some other event *V*_*K*_.

## Literature Cited

Abramowitz, M. and Stegun, I. A., editors 1972. Handbook of mathematical functions with formulas, graphs, and mathematical tables, 9th printing. Dover, New York.

Bollback, J. P., York, T. L., Nielsen, R. 2008. Estimation of 2N_e_s from temporal allele frequency data. Genetics, 179: 497–502.

Bonhoeffer, S., Barbour, A. D., and De Boer, R. J. 2002. Procedures for reliable estimation of viral fitness from time-series data. Proc. R. Soc. Lond. B., 269: 1887–1893.

Burke, M. 2012. How does adaptation sweep through the genome? Insights from long-term selection experiments. Proc. Roy. Soc. Lond. B, page rspb20120799.

Clark, A. 1979. The effects of interspecific compet0ition on the dynamics of a polymorphism in an experimental population of Drosophila melanogaster. Genetics, 92: 1315–1328.

Clarke, B. Murray, J. 1962. Changes in gene-frequency in Cepaea nemoralis (L.): the estimation of selective values. Heredity, 17: 467–476.

Cook, L. M., Cowie, R. H., Jones, J. S. 1999. Change in morph frequency in the snail Cepaea nemoralis on the Marlborough Downs. Heredity, 82: 336–342.

Cook, L. M., Sutton, S. L., Crawford, T. J. 2005. Melanic moth frequencies in Yorkshire, an old English industrial hot spot. Journal of Heredity, 96: 522–528.

Cowie, R. H. Jones, J. S. 1998. Gene frequency changes in Cepaea snails on the Marlborough Downs over 25 years. Biological journal of the Linnean Society, 65: 233–255.

Durrett, R. 2008. Probability models for DNA sequence evolution. Springer Science & Business Media.

Ewens, W. J. 1963. Numerical results and diffusion approximations in a genetic process. Biometrika, 50: 241–249.

Ewens, W. J. 2004. Mathematical Population Genetics: I, 2nd ed. Springer.

Feder, A. F., Kryazhimskiy, S., Plotkin, J. B. 2014. Identifying signatures of selection in genetic time series. Genetics, 196: 509–522.

Felsenstein, J. 1976. The theoretical population genetics of variable selection and migration. Annual Review of Genetics, 10: 253–280.

Ferrer-Admetlla, A.., Leuenberger, C., Jensen, J. D., Wegmann, D. 2015. An approximate markov model for the wright-fisher diffusion. Genetics. doi:10.1534.

Fisher, R. A. 1922. On the dominance ratio. Proceedings of the royal society of Edinburgh, 42: 321–341.

Fisher, R. A. Ford, E. B. 1947. The spread of a gene in natural conditions in a colony of the moth Panaxia dominula L. Oliver & Boyd.

Foll, M., Shim, H., Jensen, J. D. 2015. WFABC: a Wright–Fisher ABC-based approach for inferring effective population sizes and selection coefficients from time-sampled data. Molecular Ecology Resources, 15: 87–98.

Gallet, R., Cooper, T. F., Elena, S. F., and Lenormand, T. 2012. Measuring selection coefficients below 10^-3^: method, questions, and prospects. Genetics, 190(1): 175–186.

Gillespie, J. H. 2010. Population genetics: a concise guide. JHU Press.

Goudsmit, J., De Ronde, A., Ho, D. D., Perelson, A. S. 1996. Human Immunodeficiency Virus fitness in vivo: calculations based on a single zidovudine resistance mutation at codon 215 of reverse transcriptase. Journal of virology, 70: 5662–5664.

Haldane, J. B. S. 1927. A mathematical theory of natural and artificial selection, Part V: selection and mutation. Mathematical Proceedings of the Cambridge Philosophical Society, 23: 838–844.

Harrigan, P. R., Bloor, S., Larder, B. A. 1998. Relative replicative fitness of zidovudine-resistant Human Immunodeficiency Virus Type 1 isolates in vitro. Journal of Virology, 72: 3773–3778.

Hartl, D. L. Clark, A. G. 2007. Principles of Population Genetics, 4th ed. Sinauer Associates.

Haubruge, E. Arnaud, L. 2001. Fitness consequences of malathion-specific resistance in Red Flour Beetle (Coleoptera: Tenebrionidae) and selection for resistance in the absence of malathion. Journal of economic entomology, 94(2): 552–557.

Hein, J., Schierup, M. H., Wiuf, C. 2005. Gene Genealogies, Variation and Evolution. Oxford University Press, Milton Keynes, U.K.

Illingworth, C. J. R., Parts, L., Schiffels, S., Liti, G., Mustonen, V. 2012. Quantifying selection acting on a complex trait using allele frequency time series data. Mol. Biol. Evol., 29: 1187–1197.

Jenkins, P. A. Spanò, D. 2015. Exact simulation of the Wright-Fisher diffusion. arXiv:1506.06998, http://arxiv.org/abs/1506.06998.

Karlin, S. Taylor, H. 1981. A second course in stochastic processes, Second Ed. Academic Press.

Labbé, P., Sidos, N., Raymond, M., and Lenormand, T. 2009. Resistance gene replacement in the mosquito Culex pipiens: fitness estimation from long-term cline series. Genetics, 182: 303–312.

Lacerda, M. Seoighe, C. 2014. Population genetics inference for longitudinally-sampled mutants under strong selection. Genetics, 198: 1237–1250.

Long, A., Liti, G., Luptak, A., Tenaillon, O. 2015. Elucidating the molecular architecture of adaptation via evolve and resequence experiments. Nat. Rev. Genet., 16: 567–582.

Lynch, M. 1987. The consequences of fluctuating selection for isozyme polymorphisms in daphnia. Genetics, 115: 657–669.

Malaspinas, A., Malaspinas, O., Evans, S. N., Slatkin, M. 2012. Estimating allele age and selection coefficient from time-serial data. Genetics, 192: 599–607.

Manly, B. F. 1985. The statistics of natural selection. Chapman & Hall.

Mathieson, I. McVean, G. 2013. Estimating selection coefficients in spatially structured populations from time series data of allele frequencies. Genetics, 193: 973–984.

Nishino, J. 2013. Detecting selection using time-series data of allele frequencies with multiple independent reference loci. G3, 3: 2151–2161.

O’Hara, R. B. 2005. Comparing the effects of genetic drift and fluctuating selection on genotype frequency changes in the scarlet tiger moth. Proc. Roy. Soc. Lond. B, 272: 211–217.

Rabiner, L. R. 1989. A tutorial on Hidden Markov Models and selected applications in speech recognition. Proc. IEEE, 77: 257–286.

Reimchen, T. E. Nosil, P. 2002. Temporal variation in divergent selection on spine number in Threespine Stickleback. Evolution, 56: 2472–2483.

Rouzine, I. M., Rodrigo, A., Coffin, J. M. 2001. Transition between stochastic evolution and deterministic evolution in the presence of selection: general theory and application to virology. Microbiology and molecular biology reviews, 65: 151–185.

Schaffer, H. E., Yardley, D., Anderson, W. W. 1977. Drift or selection: a statistical test of gene frequency variation over generations. Genetics, 87: 371–379.

Siepielski, A. M., DiBattista, J. D., Carlson, S. M. 2009. Its about time: the temporal dynamics of phenotypic selection in the wild. Ecology Letters, 12(11): 1261–1276.

Song, Y. S. and Steinrücken, M. 2012. A simple method for finding explicit analytic transition densities of diffusion processes with general diploid selection. Genetics, 190(3): 1117–1129.

Steinrücken, M., Bhaskar, A., Song, Y. S. 2014. A novel spectral method for inferring general diploid selection from time series genetic data. Annals of Applied Statistics, 8(4):2203–2222.

Steinrücken, M., Jewett, E. M., Song, Y. S. 2015. Spectraltdf: transition densities of diffusion processes with time-varying selection parameters, mutation rates and effective population sizes. Bioinformatics, page btv627.

Stine, O. C. Smith, K. D. 1990. The estimation of selection coefficients in afrikaners: Huntington disease, porphyria variegata, and lipoid proteinosis. Am. J. Hum. Genet., 46:452–458.

Topa, H., Jónás, A., Kofler, R., Kosiol, C., Honkela, A. 2015. Gaussian process test for high-throughput sequencing time series: application to experimental evolution. Bioinformatics, 31: 1762–1770.

Wakeley, J. 2008. Coalescent theory: An introduction. Roberts & Company Publishers, Greenwood Village, CO.

Wall, S., Carter, M. A., Clarke, B. 1980. Temporal changes of gene frequencies in cepaea hortensis. Biological Journal of the Linnean Society, 14(3-4): 303–317.

Wilson, S. R. 1980. Analyzing gene-frequency data when the effective population size is finite. Genetics, 95: 489–502.

Zhao, L., Lascoux, M., Waxman, D. 2014. Exact simulation of conditioned wright–fisher models. Journal of theoretical biology, 363: 419–426.

